# Computational design of non-porous, pH-responsive antibody nanoparticles

**DOI:** 10.1101/2023.04.17.537263

**Authors:** Erin C. Yang, Robby Divine, Marcos C. Miranda, Andrew J. Borst, Will Sheffler, Jason Z Zhang, Justin Decarreau, Amijai Saragovi, Mohamad Abedi, Nicolas Goldbach, Maggie Ahlrichs, Craig Dobbins, Alexis Hand, Suna Cheng, Mila Lamb, Paul M. Levine, Sidney Chan, Rebecca Skotheim, Jorge Fallas, George Ueda, Joshua Lubner, Masaharu Somiya, Alena Khmelinskaia, Neil P. King, David Baker

## Abstract

Programming protein nanomaterials to respond to changes in environmental conditions is a current challenge for protein design and important for targeted delivery of biologics. We describe the design of octahedral non-porous nanoparticles with the three symmetry axes (four-fold, three-fold, and two-fold) occupied by three distinct protein homooligomers: a *de novo* designed tetramer, an antibody of interest, and a designed trimer programmed to disassemble below a tunable pH transition point. The nanoparticles assemble cooperatively from independently purified components, and a cryo-EM density map reveals that the structure is very close to the computational design model. The designed nanoparticles can package a variety of molecular payloads, are endocytosed following antibody-mediated targeting of cell surface receptors, and undergo tunable pH-dependent disassembly at pH values ranging between to 5.9-6.7. To our knowledge, these are the first designed nanoparticles with more than two structural components and with finely tunable environmental sensitivity, and they provide new routes to antibody-directed targeted delivery.

## Main Text

There is considerable interest in tailoring nanoparticle platforms for targeted delivery of therapeutic molecules. Effective targeted delivery nanoparticle platforms require assembly and encapsulation of molecules outside the target cell, followed by target recognition, triggered nanoparticle disassembly, and controlled cargo release once inside the cell ^1–9^. Cellular uptake of extracellular molecules such as protein-based nanoparticles via endocytosis involves traversal of various membrane-bound organelles, including the low-pH endosome and lysosome ^10–13^. While a number of self-assembling protein nanoparticles with customized structures have been designed, they are composed of just one or two unique, static building blocks and efforts to adapt them for cargo packaging and delivery applications are still in their infancy ^1,14–20^. Particularly attractive for delivery applications are antibody-incorporating nanoparticles, where one component is a designed homooligomer that, upon mixing with any antibody of interest, generates a bounded, multivalent, polyhedral assembly ^21^ (**Fig 1A**). While such antibody nanoparticles can enhance cell signaling activity and have demonstrated a broad utility for selectively targeting cell surface receptors, they are quite porous, which complicates the packaging and retention of molecular cargoes.

**Figure 1:**
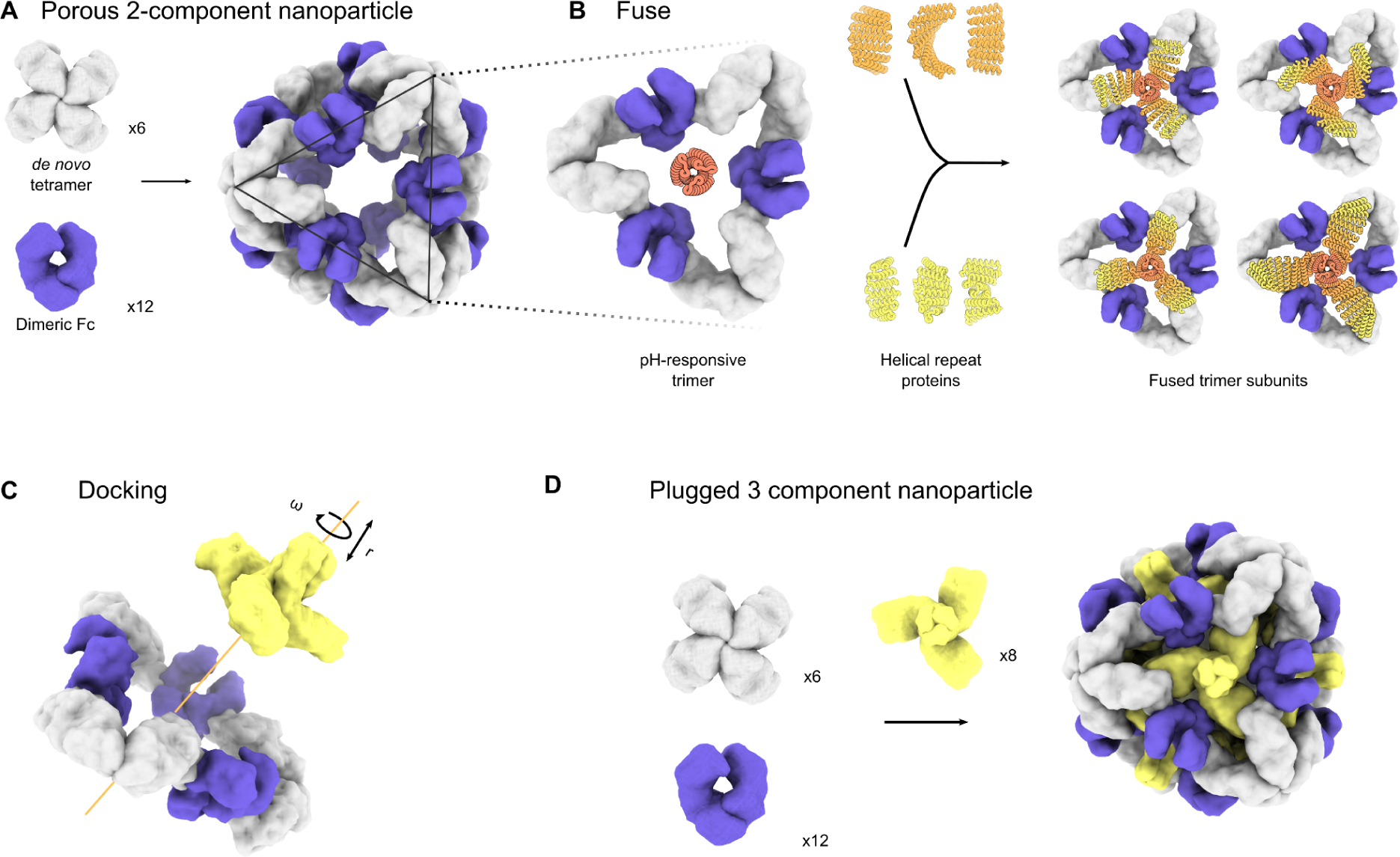
Design of symmetry-matched plugs to fill empty symmetry axes in protein nanoparticles. **A.** 6 *de novo* tetramers (gray) and 12 dimeric Fc domains (purple) assemble into a porous octahedral O42 nanoparticle. The tetramers are aligned along the 4-fold symmetry axis and the Fc domains along the 2-fold symmetry axis. **B.** Combinations of helical repeat proteins were fused to each other and to the pH trimer subunit at regions of high backbone overlap between pairs of helices to generate fused trimer subunits large enough to fully occupy the void along the 3-fold axis in the original nanoparticle. **C.** These pH-dependent trimers were then docked into the nanoparticle by rotating and translating along its 3-fold axis. **D.** The resulting three-component nanoparticle has 8 new trimeric subunits (yellow) which occupy the three-fold symmetry axis of the octahedral architecture.

To enable packaging and pH-dependent release of molecular cargoes, we sought to close off the apertures in the previously designed antibody nanoparticles with the addition of computationally designed pH-responsive proteins. We focused on octahedral antibody nanoparticles (O42.1) constructed from a C4-symmetric designed tetramer and C2-symmetric IgG dimers, which are aligned along the C4 and C2 symmetry axes of the octahedral architecture (**Fig 1A**). On the open C3 axis, we aimed to incorporate a designed pH-dependent C3 trimer ^22^ and tune it to disassemble at pH values corresponding to the native environment of the endosome (**Fig 1B**). We reasoned that such three-component nanoparticles could 1) selectively enter target cells, 2) encapsulate molecular cargoes without leakage, and 3) disassemble in the acidic environment of the endosome.

The previously designed pH-dependent trimer is much smaller than the aperture along the C3 axis of the octahedral nanoparticle (**Fig 1B**), and hence filling the C3 axis of the octahedral nanoparticles with the pH-dependent trimer required extending the backbone of the trimer such that it could contact and make shape-complementary interactions with the designed tetramer. To enable this, we combined helical fusion ^23,24^ and protein docking ^25^ approaches into a single design pipeline. We extended the pH trimer by fusing combinations of helical repeat protein building blocks onto each subunit to generate a total of 80,000 distinct C3 fusions with helical repeats of variable geometry extending outwards from the C3 axis (**Fig 1B**, see methods). The resultant diverse set of C3 building blocks were docked into the three-fold pore by aligning both C3 axes and sampling translational and rotational degrees of freedom along this axis ^25^ (**Fig 1C**). Variable-length truncations of the terminal helices of the repeat protein arms were evaluated to optimize the docked interface between the new C3 building block and the C4 subunits of the octahedral assembly (see methods). The resulting “plugged” octahedral assembly (O432) contains twelve IgG1-Fc domains along the octahedral two-fold axes, six tetramers along the octahedral four-fold axes, and eight trimeric plugs along the octahedral three-fold axes (**Fig 1D**).

The newly generated interfaces between the pH trimer fusions and the octahedral assembly were evaluated for designability using a combination of the residue pair transform (rpx) score—a prediction of interaction energy following sequence design ^25,26^—and overall shape complementarity at the interface ^27^. For 6000 docks that were predicted to have high designability and shape complementarity, the amino acid sequence at the newly formed fusion junctions and at the interface between the trimer and antibody nanoparticle were optimized using Rosetta sequence design calculations ^28^. This design step introduced mutations on both the trimer and the tetramer subunits. Designed interfaces were evaluated for secondary structure contacts and chemical complementarity, and 45 designed trimeric plug and nanoparticle tetramer pairs were selected for experimental characterization. The designed interfaces intentionally spanned a broad range of buried solvent accessible surface area (SASA; 1500–3000 Å^2^), as the optimal interface size for this type of symmetric assembly was unknown.

Designed trimers and tetramers with C-terminal 6×-histidine tags on the tetramer were expressed bicistronically in *E. coli* and subject to immobilized metal affinity chromatography (IMAC) purification. SDS-polyacrylamide gel electrophoresis (SDS-PAGE) identified 16 of 45 designs where the designed trimer co-eluted with the tetramer. A sfGFP-Fc fusion protein ^21^ was added to the clarified lysates of these co-expressed trimers and tetramers, and the resulting assemblies were purified via IMAC. SDS-PAGE and native PAGE were used to determine which designs formed three-component assemblies (**Fig S1A-B**). For the 5 out of 16 designs in which all three components associated, genes expressing the trimer and tetramer were subcloned into separate expression vectors, and the independently expressed oligomers were purified separately by size exclusion chromatography (SEC) on a Superdex 200 10/300 GL column (**Fig S1C**). Three-component assemblies were prepared by mixing together the three purified proteins—trimeric plug, tetramer, and the Fc of human IgG1—in a 1:1:1 stoichiometric ratio, followed by overnight incubation at 25°C and purification by SEC using a Superdex 6 10/300 GL column (**Fig S1D**). Due to tetramer insolubility, insufficient material was produced to prepare a three-component assembly reaction for one design, O432-43.

Since the addition of the trimeric plug should not affect the diameter of the octahedral antibody nanoparticle, we next compared the SEC elution volumes of all three-component mixtures to the elution profile of the original O42.1 antibody nanoparticle ^21^. Mixing all three components in a 1:1:1 stoichiometric ratio yielded elution peaks in the void volume with shoulders between 9-11 mL (**Fig S1D**), where the shoulder peaks matched the elution profile of the original O42.1 antibody nanoparticle ^21^ (**Fig S1E**). Non-reducing SDS-PAGE of both the void and shoulder peaks showed that only one assembly (O432-17) contained all three protein components in the elution fraction (**Fig S1F-G**). Optimization of the stoichiometric ratios of each protomer present during *in vitro* assembly resulted in the O432-17 peak shifting from the void volume toward the expected elution volume during SEC, which matched the profile of the original O42.1 antibody nanoparticle (**Fig 2B**). The optimal assembly ratio per protomer of purified trimeric plug, tetramer, and Fc was determined to be 1.1:1.1:1. Dynamic light scattering (DLS) of the main O432 peak indicated a hydrodynamic diameter of 34 nm and a polydispersity index (PDI) of 0.05 (**Fig 2C**) and negative-stain electron microscopy (NS-EM) revealed monodisperse nanoparticles (**Fig 2D**). Two-dimensional class averages of negatively stained micrographs revealed plug-like density in the 3-fold views compared to the original two-component antibody nanoparticle.

**Figure 2:**
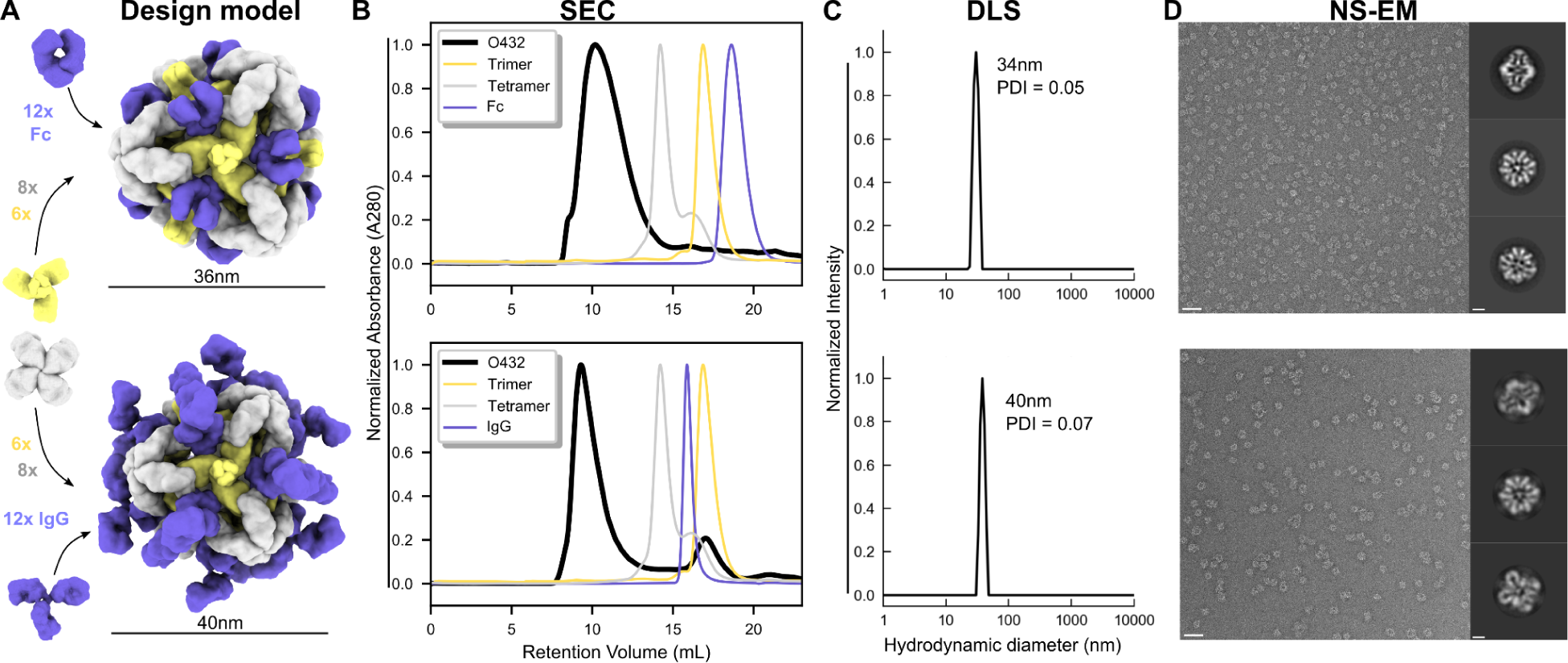
Mixing independently purified components enables stable, efficient assembly with both Fc and IgG. **A.** Design models with Fc or IgG (purple), designed nanoparticle-forming tetramers (gray), and pH-dependent plug-forming trimers (yellow). **B.** Overlay of representative SEC traces of full assembly formed by mixing designed tetramers, trimers, and Fc or IgG (black) with those of the single components in gray (tetramer), yellow (trimeric plug), or purple (Fc or IgG). **C.** Representative DLS of fractions collected from the O432 assembly peak shows average hydrodynamic diameters of 34 nm (polydispersity index or PDI: 0.05) and 40 nm (PDI: 0.05) for the O432 assemblies with Fc and full-length IgG, respectively. **D.** Negatively stained electron micrographs with reference-free 2D class averages along each axis of symmetry in inset; electron microscopy images were collected prior to SEC purification. Scale bars, 100 nm and 10 nm for the micrograph and 2D averages, respectively.

We found that assembly of the designed nanoparticle was cooperative and required all three components: mixing any two of the three O432-17 components stoichiometrically did not result in nanoparticles (**Fig S2A**). As expected, mixing the trimeric plug and Fc resulted in no assembly, as there is no designed interface between those two components. Mixing the tetramer with Fc resulted in visible aggregate and partially formed nanoparticles (**Fig S2B**), likely due to the formation of off-target interactions by the newly designed hydrophobic plug interface. Mixing the tetramer and trimeric plug did not result in association even at high concentration (400 µM per monomer) (**Fig S2C**). This cooperativity simplifies preparation of the three component nanoparticles as it prevents incomplete assembly of nanoparticles containing two out of the three components, and thus eliminates the need for additional purification steps to separate these species from the intended three-component assembly.

For downstream delivery applications, we tested whether the O432-17 nanoparticle would assemble when the designed trimer and tetramer were co-incubated with full-length IgG antibodies containing both Fc and Fab domains (**Fig 2B-D**). The O432-17 design eluted in the void volume, owing to the increased diameter from the additional Fab domains (**Fig 2B**). DLS of this void volume peak revealed a monodisperse hydrodynamic diameter of 40 nm and PDI of 0.07, in line with the expected diameter of the IgG-containing O432-17 assembly (**Fig 2C**). NS-EM micrographs and two-dimensional class averages of this peak fraction exhibited plug-like density in three-fold views following 2D classification as well as Fab-like density in two-fold, three-fold, and four-fold views (**Fig 2D**). Despite the clear presence of additional density corresponding to the IgG Fab domains, due to the inherent marked flexibility between the IgG Fc and Fab domains ^29^, the Fab domains are largely averaged out in the 2D averages of this complex (**Fig 2D**).

We next sought to determine the completeness of the three-component Fc-containing octahedral nanoparticle assembly and the accuracy of the newly designed interface between the trimer and tetramer using single-particle cryo-EM (**Fig 3A-D, Fig S3E**). Following data collection and preprocessing of raw micrographs, a subset of the best selected two-dimensional averages with fully assembled O432-17 nanoparticles (**Fig S3A-B**) were used to generate an *ab initio* 3D reconstruction in the absence of applied symmetry (**Fig S3C**). This initial map and the corresponding set of particles were next subclassified into four distinct classes following 3D heterogeneous refinement (**Fig S3D**), also in the absence of any applied symmetry operator. All four subclasses revealed the presence of fully plugged antibody nanoparticles, demonstrating complete O432 assembly for this system at the concentrations used. We observe density occupying all symmetry axes of the octahedral architecture with six designed tetramers at the four-fold octahedral symmetry axes forming interfaces with eight designed trimers at the three-fold symmetry axes and twelve dimeric Fc fragments at the two-fold symmetry axis. Upon confirming the completeness of this nanoparticle assembly, a subsequent 3D refinement containing particles from all four of the aforementioned classes was generated after applying octahedral symmetry, resulting in a final map with an estimated global resolution of 7 Å (**Fig S3C**).

**Figure 3:**
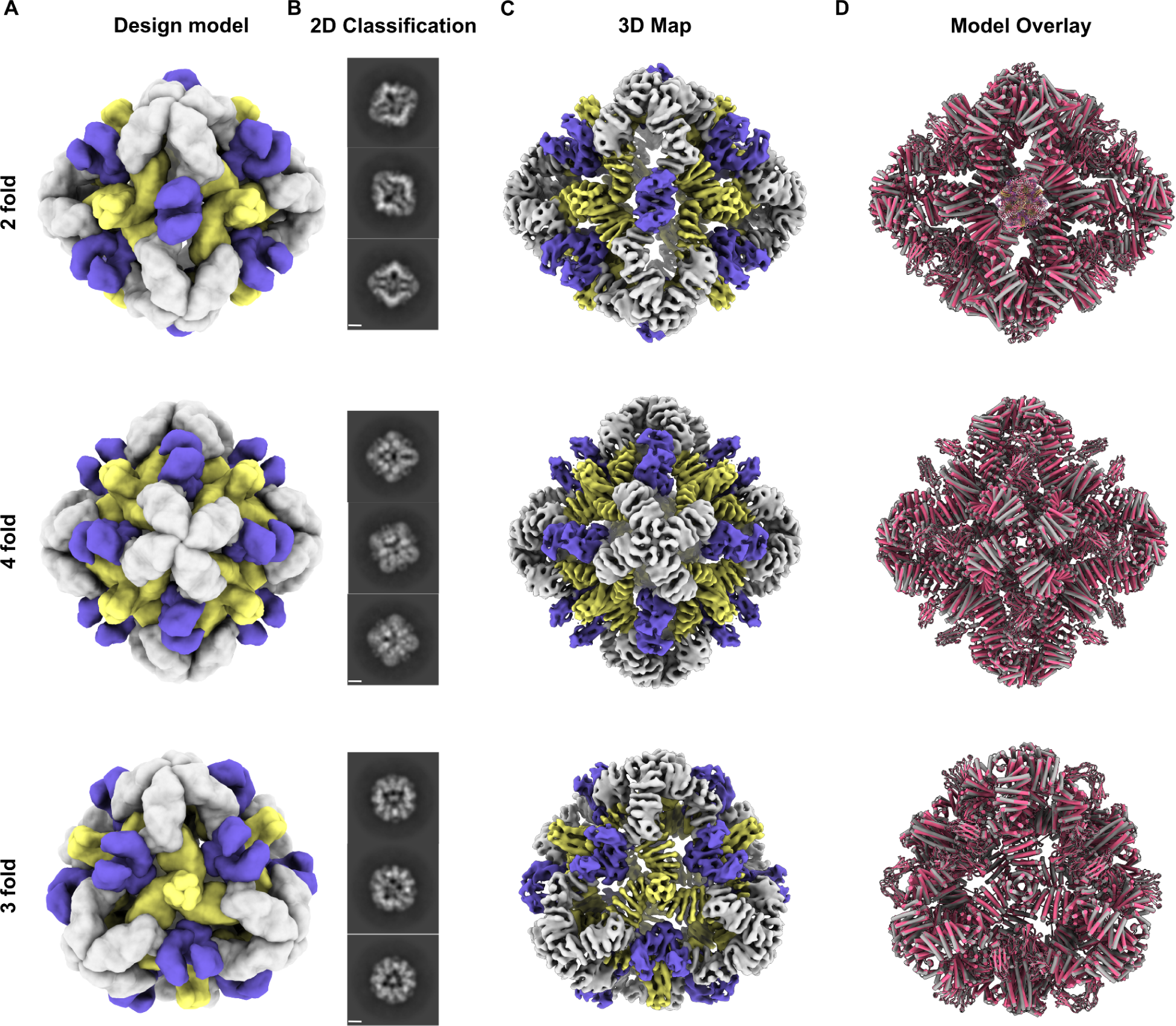
Cryo-EM analysis of 72-subunit nanoparticles composed of three distinct structural components. **A.** The O432 assembly with Fc before SEC purification was characterized by cryogenic electron microscopy. Computational design models viewed along each axis of symmetry of the octahedral architecture are shown. **B.** Representative 2D class averages along each axis of symmetry. Scale bar, 10 nm. **C.** 7 Å 3D EM density map reconstructed from the collected dataset. **D.** Overlay of the design model (gray) and the design model relaxed into the 3D reconstruction (pink) showing high agreement. Encouraged by these structural results, we next set out to redesign the nanoparticles to have either highly positively or highly negatively charged interiors to enable packaging of molecular cargoes via electrostatic interactions ^17,30,31^. We focused our redesign on selected interior surface residues of the trimeric plug, favorably weighting mutations to amino acids with the desired charge (or no charge), and unfavorably weighting mutations to amino acids with the undesired charge. We screened for packaging of positively charged GFP (pos36GFP) ^32^ by mixing negatively charged trimeric plug variants exhibiting different magnitudes of interior surface charge with tetramer, Fc, and pos36GFP (**Fig 4A**). One variant interacted with supercharged GFP in these conditions, as shown by co-elution of the 488 nm signal and the 280 nm signal during SEC (**Fig 4B**), and will be referred to subsequently as O432-17(-).

We estimated the accuracy of our design protocol by relaxing the original computational O432-17 design model in the experimentally determined cryo-EM map. The relaxed model of O432-17 containing the plug, tetramer, and Fc is in close agreement with the original design (**Fig 3D**). We observed cryo-EM density corresponding to the helices of the tetramers at the four-fold vertices, where each helical repeat extends from the four-fold axis along the edges of the octahedral architecture to bind Fc. The cryo-EM density also shows the helices of the trimeric components along the faces of the octahedral architecture, where the helical repeats extend to form an interface with the tetramers at the edges. The addition of the trimeric plug reduces porosity of the antibody nanoparticle as intended: the largest diameter pore is 3 nm compared to 13 nm in the original O42.1 nanoparticle ^21^. While there is some structural distortion in the helical repeat regions of the trimeric plug and tetramer, suggesting slight structural flexibility of the helical repeat domains or fusions between helical repeat regions, the helices forming the trimer and tetramer interface are largely consistent between the design and relaxed model (**Fig S3F**). The Fc domain exhibited very little secondary structure deviation between the design model and relaxed model.

Comparing the relaxed model of our plugged O432-17 nanoparticle to that of the original O42.1 antibody nanoparticle design model ^21^ showed that the relaxed cryo-EM model is significantly closer to the original O42.1 design model than the previously determined O42.1 cryo-EM structure. The Cα RMSDs of the asymmetric units are 1.6 Å between the O432-17 relaxed model and its design model, 1.9 Å between the O432-17 design model and O42.1 relaxed model, and 4.2 Å between the the O42.1 relaxed model and its design model. These results suggest that the addition of the trimeric plug buttresses the tetramer into a conformation that more closely matches the original design model, and may also reduce the overall flexibility of the system, as compared to the original O42.1 design (**Fig S4A-C**).

Including 1 M NaCl in the buffers used during packaging and SEC prevented packaging (**Fig 4C**), suggesting that GFP packaging was largely driven by electrostatic interactions between the cargo and nanoparticle interior. Trimeric plug variants of varying interior positive charge magnitudes were then screened for packaging of a 154-nt prime editing guide RNA (pegRNA) cargo ^33–35^. We mixed the positively charged trimer variants, tetramer, and the α-EGFR antibody Cetuximab (CTX) with the pegRNA (**Fig 4A**), carried out non-denaturing electrophoresis, and stained with SYBR Gold (RNA) and Coomassie (protein) with and without Benzonase treatment. For one variant, which will be referred to as O432-17(+), comigration of nucleic acid and protein with and without Benzonase treatment indicated successful packaging of nucleic acid (**Fig 4D-E**). Excess nucleic acid that did not comigrate with the protein was degraded in the Benzonase-treated sample, demonstrating nanoparticle protection of the nucleic acid that comigrated with O432-17(+).

**Figure 4:**
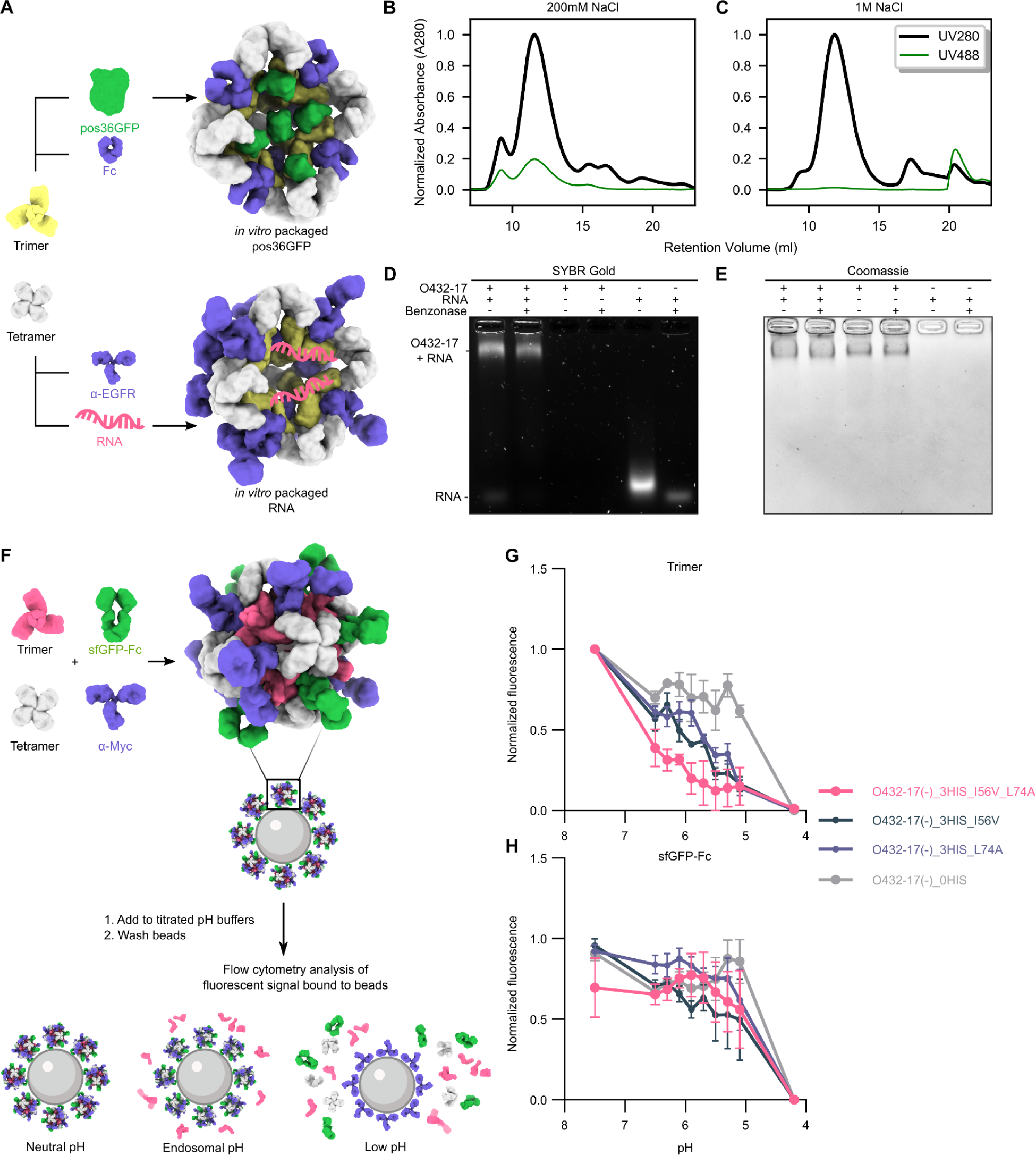
Plugged antibody nanoparticles electrostatically package cargoes and disassemble in response to acidification. **A.** Positively or negatively charged variants of the designed trimers and designed tetramers were either assembled with pos36GFP and human Fc (top) or RNA and ⍺-EGFR mAb (bottom). **B.** SEC chromatograms of *in vitro* packaging reactions with O432-17(-) were performed in either 200 mM NaCl or **C.** 1 M NaCl. Absorbance was monitored at 280 nm (black) and 488 nm (green). **D.** O432-17(+) assembled *in vitro* with RNA was treated with Benzonase, electrophoresed on non-denaturing 0.8% agarose gels, and stained with SYBR gold (nucleic acid) and **E.** Coomassie (protein). **F.** pH titration experimental design. O432-17 nanoparticles were assembled with AF647-conjugated trimeric plug variants, designed tetramer, sfGFP-Fc, and α-Myc antibody. Nanoparticles were incubated with Myc peptide-coated beads and split into titrated pH buffers. Beads were washed in TBS pH 7.5 and the remaining trimer and sfGFP-Fc fluorescence remaining on the beads was analyzed by flow cytometry. **G.** AF647 and **H.** sfGFP fluorescence were normalized to the minimum and maximum values across the titration and analyzed as a function of pH for each nanoparticle variant.

NS-EM micrographs and two-dimensional class averages confirmed three-component nanoparticle assembly of O432-17(+) in the presence of RNA (**Fig S5A-B**). Thus, our designed three-component nanoparticles efficiently encapsulate and protect molecular cargoes.

We next explored whether O432-17(-) and O432-17(+) disassembled below pH 6.0 using a medium-throughput, low-concentration flow cytometry-based assay. We conjugated AlexaFluor647 (AF647) via maleimide chemistry to a C-terminal cysteine on the O432-17(-) trimeric plug variant. We assembled O432-17 nanoparticles with the tetramer, AF647-conjugated trimeric plug, and a 50/50 mixture of α-myc mouse mAb (Cell Signaling Technologies 9B11) and sfGFP-Fc, the latter of which acted as a marker for a non-plug nanoparticle component. Assembled nanoparticles were incubated with 3 μm polystyrene beads coated with biotinylated myc-tag for one hour to allow efficient loading of the α-myc containing nanoparticles onto the beads. The beads were then split into an excess volume of titrated citrate-phosphate buffers ranging from pH 4.2 to pH 7.5 and were incubated at 25°C for 30 minutes. After 30 minutes, the beads were resuspended and brought back to pH 7.5 to normalize sfGFP fluorescence, and then analyzed for AF647 and sfGFP signal by flow cytometry (**Fig 4F**). After isolating singlet beads, we normalized the mean fluorescence intensity of the AF647 on the trimer and sfGFP on the Fc at each pH (**Fig S6A-D**) to the minimum and maximum values observed across the pH titration. The O432-17(-) plug design (named O432-17(-)_2HIS), which was based on a pH-responsive helical bundle with two histidine hydrogen bond networks that dissociated into monomers at pH 4.9 ^22^, did not show reduction in AF647 or sfGFP fluorescence until below pH 5.1, similar to a negative control trimer variant with all histidines substituted with asparagine (named O432-17(-)_0HIS; **Fig S6E-F**). This result indicated that the sensitivity of the trimeric plug to pH was dampened relative to the original pH-responsive helical bundle, likely due to stabilization afforded by nanoparticle assembly ^36^.

To improve the pH sensitivity of the O432-17(-) trimeric plug design, we introduced a third histidine hydrogen bond network (3HIS) and point mutations at two residues involved in hydrophobic packing interactions at the trimeric interface within each protomer, I56V and L74A (named O432-17(-)_3HIS_I56V and O432-17(-)_3HIS_L74A). O432-17(-)_3HIS_I56V and O432-17(-)_3HIS_L74A each showed clear pH-dependent release of the pH trimer (**Fig 4G**). The apparent pKas of trimer dissociation (the pH where the AF647 fluorescence was 50% of the maximum signal) were pH 6.1 and pH 5.9 for O432-17(-)_3HIS_I56V and O432-17(-)_3HIS_L74A, respectively (**Fig S6E**). The apparent pKa of sfGFP fluorescence for these variants also shifted to more basic pH (pH 5.3 and pH 4.7, respectively) relative to the negative control trimer variant (named O432-17(-) 0 HIS networks), but less so than the AF647 apparent pKa, suggesting that nanoparticle disassembly was less pH-sensitive than plug dissociation from the nanoparticle (**Fig S6F**). O432-17(-)_3HIS_I56V_L74A, containing both point mutations per protomer, had an AF647-based apparent pKa of pH 6.7 and sfGFP-based apparent pKa of pH 5, indicating a synergistic effect of the two point mutations (**Fig 4G-H**). These pH-responsive, O432-17(-) variants did not encapsulate pos36GFP, suggesting that weakening the trimeric interface with an additional histidine hydrogen-bond network and up to two hydrophobic packing mutations per protomer reduces the assembly efficiency of the fully plugged nanoparticle in the presence of molecular cargo, even when the histidine network and point mutations were not designed at the interface between the trimeric plug and the tetramer.

We also introduced a third histidine hydrogen-bond network and the same two hydrophobic packing mutations in the O432-17(+) trimeric plug design. We generated four variants, two containing the third histidine hydrogen-bond network and either the I56V or L74A point mutation (O432-17(+)_3HIS_I56V and O432-17(+)_3HIS_L74A), one containing the third histidine hydrogen-bond network and both I56V and L74A point mutations (O432-17(+)_3HIS_ I56V_L74A), and one negative control containing zero histidine hydrogen bond networks (O432-17_0HIS), where all histidines were mutated to asparagine. We compared the apparent pKa, calculated from the AF647 and sfGFP fluorescence signals, of each positively charged plug variant over the pH titration relative to the negative control. All pH-responsive O432-17(+) trimeric plug variants showed an apparent pKa of pH 6.1 for AF647 and pH 5.8 for sfGFP fluorescence, respectively. Unlike the negatively charged variants, we did not observe a synergistic effect when combining the I56V and L74A mutations within the positively charged plug variants (**Fig S6G-H**). The apparent pKa of AF647 fluorescence for O432-17(+) variants containing histidine hydrogen-bond networks was more basic than the negative control (**Fig S6G-H**). The apparent pKa of sfGFP fluorescence for each O432-17(+) trimeric plug variant containing three histidine hydrogen-bond networks was only slightly more acidic than the apparent pKa for AF647 fluorescence, suggesting that nanoparticle dissociation and plug dissociation from the nanoparticle were equally pH-sensitive for the O432-17(+) nanoparticles.

We next tested the ability of the O432-17 nanoparticles to enter cells through receptor-mediated endocytosis. We assembled targeted nanoparticles by incubating a 1:1 stoichiometric mixture of CTX and Fc fused to mRuby2 (mRuby2-Fc) with the tetramer and a pH-responsive trimeric plug variant labeled with AF647 (named O432-17-CTX). As negative controls, we assembled a non-EGFR targeting nanoparticle by mixing the tetramer and trimer-AF647 with mRuby2-Fc (**Fig 5A**) (named O432-17-Fc). Assembled O432-17-CTX and O432-17-Fc were incubated at a final concentration of 10 nM per monomer in serum-free media for up to 16 hours with either A431 (high EGFR expression) wild-type (WT), HeLa WT (moderate EGFR expression), or HeLa EGFR knockout (KO) cells. After three hours of nanoparticle treatment, cells were fixed and immunostained with LAMP2A (late endosome and lysosomal marker) and imaged with confocal microscopy. We observed mRuby2 and AF647 fluorescence in A431 cells incubated with O432-17-CTX but little to no mRuby2 and AF647 fluorescence in cells incubated with O432-17-Fc, suggesting that the O432-17(-) nanoparticles can enter cells via EGFR-dependent endocytosis (**Fig 5B**). We also observed mRuby2 and AF647 fluorescence in HeLa WT cells incubated with O432-17-CTX (**Fig S7**). O432-17-CTX nanoparticles target specifically to EGFR-expressing cells, as pH-dependent O432-17-CTX nanoparticles did not accumulate in HeLa EGFR KO cells (**Fig S7**).

**Figure 5:**
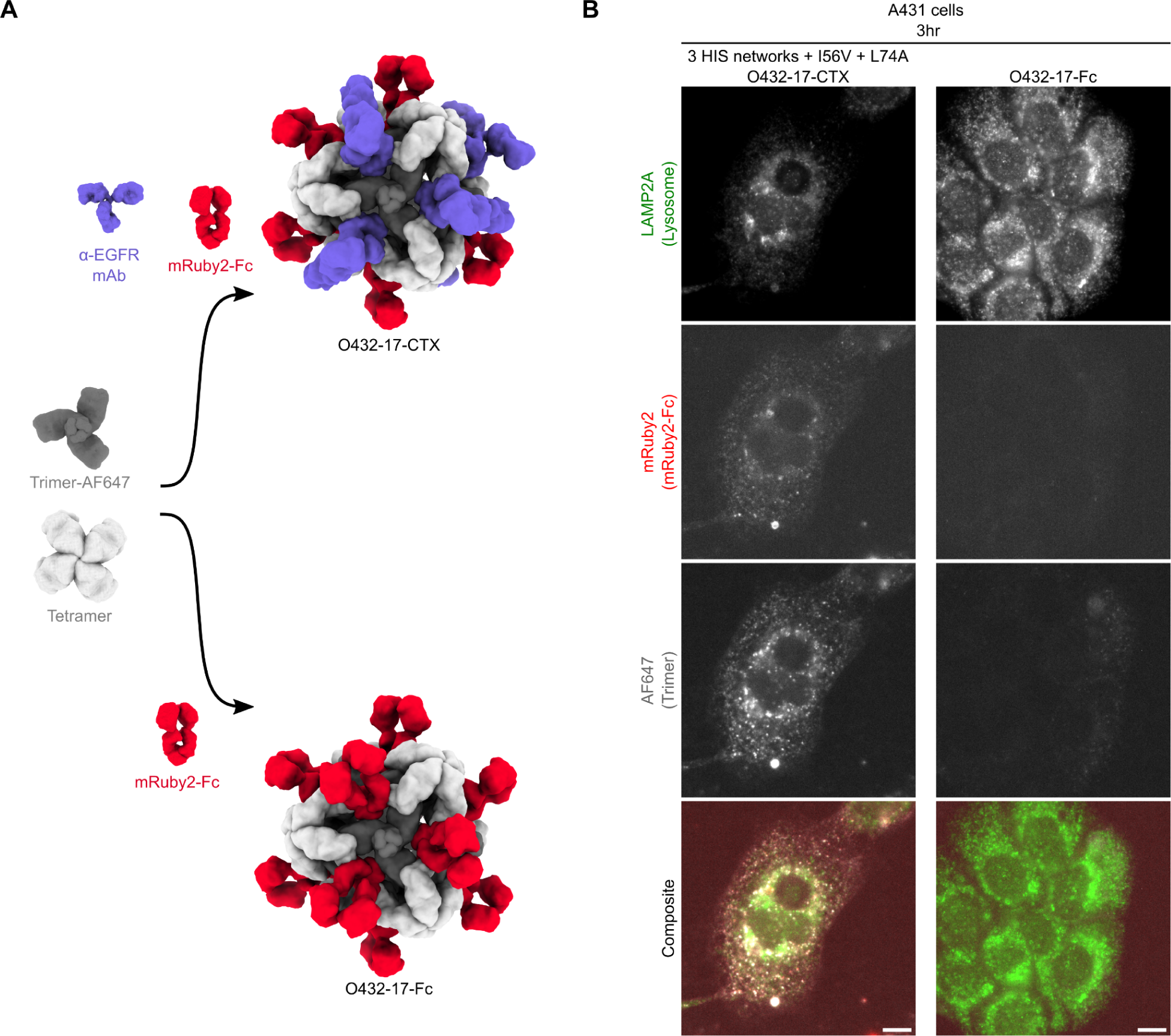
Targeted receptor-mediated uptake of O432 nanoparticles. **A.** Targeted and non-targeted variants of the O432-17(-) nanoparticle were assembled *in vitro* with trimeric plug conjugated with AF647, designed tetramer, and either premixed mRuby2-Fc and ⍺-EGFR mAb (top) or mRuby2-Fc alone (bottom). **B.** Cellular uptake of O432-17-CTX and O432-17-Fc nanoparticles was measured by the AF647 and mRuby2 fluorescence within the cell area, identified by lysosome membrane immunostaining using LAMP2A antibodies, after 3 hours of incubation in A431 cells. Images correspond to single confocal planes, and grayscale panels correspond to each channel of the composite image showing lysosomal membranes (green), mRuby2-Fc (red), and Trimer-AF647 (gray). Scale bar, 10 µm.

## Discussion

We describe a general approach for reducing the porosity of protein nanomaterials by designing custom symmetric plugs that fill pores present along unoccupied symmetry axes. Using this approach, we generated the first designed protein nanoparticles with distinct structural components on three different symmetry axes. These designed nanoparticles can modularly package molecular cargoes, undergo pH-dependent disassembly, and selectively enter cells in a programmable, antibody dependent manner. In contrast to functionalized homomeric and designed two-component protein nanoparticles ^1,8,14,17,18,21,30,37–40^, each of the three components in our designs has a specific functional role. The trimer is responsible for pH-responsive disassembly and cargo packaging, the antibody provides cell targeting functionality, and the tetramer drives assembly of the three-component nanoparticle by making an interface with both the trimer and Fc domain. This division of labor makes our designed system highly modular: targeting specificity can be altered simply by switching the antibody component to target cells of interest, and the pH of disassembly and cargo packaging specificity can be programmed by choosing the appropriate trimer. Furthermore, assembly of the nanoparticles *in vitro* from independently purified components enables facile inclusion of multiple variants of each component, such as multiple distinct antibodies.

Induced nanoparticle disassembly at biologically relevant pH is a critical challenge for engineering drug delivery platforms, and our results represent a test of our understanding of protein disassembly and assembly dynamics ^41–44^. We were able to finely tune the pH of disassembly of our O432 system through a combination of histidine hydrogen bonding networks and cavity-introducing mutations that weaken hydrophobic interactions at the trimeric component’s oligomeric interface. Through this approach, we are able to raise the apparent pKa of disassembly to a remarkably high pH of 6.7, well above the pH of the endosome and in the range of many tumor microenvironments, and close to the maximum value achievable given the pKa of histidine.

Our O432 nanoparticle system is capable of packaging and protecting both protein and nucleic acid cargoes, disassembles at biologically relevant pH with precise tunability, incorporates a wide variety of targeting moieties, and is readily internalized by target cells. The tunable pH dependence makes this system a particularly attractive platform for engineering release and delivery of drugs during early stages of endosomal maturation. However, to be a broadly useful intracellular biologics delivery system, it will be necessary to incorporate endosomal escape machinery in future designs. The nanoparticles also provide a route to conditional delivery of drugs into the tumor microenvironment. Tumor-killing or -modulating cargoes could be packaged within the nanoparticles and directed to the tumor through targeting with tumor-specific antibodies; the pH-dependent release of cargo could minimize off-tumor toxicity and systemic exposure as compared to classic direct antibody conjugation approaches by providing an additional checkpoint on proper localization. The pH-dependent disassembly, programmability, and versatility of the O432 platform provides multiple exciting paths forward for biologics delivery.

## Data Availability

All images and data were generated and analyzed by the authors, and will be made available by the corresponding authors (D.B., N.P.K.) upon reasonable request. Density maps have been deposited in the Electron Microscopy Data Bank under the accession number EMD-29602. Source data are provided with this manuscript.

## Code Availability

Source code for the fusion, docking, and design of non-porous pH-responsive antibody nanoparticles is made available at https://github.com/erincyang/plug_design. The protocol requires compilation of the worms and rpxdock repositories, which have been made available at https://github.com/willsheffler/worms and https://github.com/willsheffler/rpxdock, respectively. Source code for generating the figures in this manuscript were written by the authors and provided in the supplementary materials.

## Acknowledgments

We would like to thank James Lazarovitz, William L. White, Scott Boyken, Christian Richardson, Issa Yousif, Hojun Choi, Samuel J. Hendel, and Audrey Olshefsky for helpful discussions, Lauren Carter and Marcos Miranda for help with SEC-MALS, Yang Hsia for initial help with purifying sfGFP-Fc fusions, Michael Murphy and Cassie Ogohara for providing Fc-fusion and IgG proteins, and Mengyu Wu and Gabriella Reggiano for help with electron microscopy. We’d also like to thank Lauren Carter, Kandise Van Wormer, Ratika Krishnamurty, Kristina Herrera, Luki Goldschmidt, and Lance Stewart for general operations. This research was funded by the NSF Grant CHE-1629214 (N.P.K. and D.B.), Defense Threat Reduction Agency Grants HDTRA1-18-1-0001 and HDTRA1-19-1-0003 (N.P.K. and D.B.), the grant DE-SC0018940 funded by the U.S. Department of Energy, Office of Science (D.B.), National Institutes of Health’s National Institute on Aging grants R01AG063845 and 1R01CA240339 (N.P.K. and D.B.), the Bill and Melinda Gates Foundation #INV-010680 (N.P.K. and D.B.), the Audacious Project at the Institute for Protein Design, Washington Research Foundation and Translational Research Fund, National Science Foundation Graduate Research Fellowship program under the grant number DGE-1762114 (E.C.Y), the Helen Hay Whitney Foundation (J.Z.Z.), the DAAD PROMOS program (N.G.), and the Howard Hughes Medical Institute (D.B.).

