## Supplemental_Information for "Computational design of non-porous, pH-responsive antibody nanoparticles"

### Methods

#### Extension of pH-responsive trimeric building blocks

We used the WORMS helical fusion software [1] to generate fusions between the base pH-trimer helical bundle and pairs of 82 helical repeat proteins [2], designed to cover a broad range of axial and radial displacement between repeat units. Scaffold information was prepared in a json file, which is used as an input parameter in the WORMS software through the `--dbfiles` option. For each scaffold, the pdb file path (“`file`”), a geometric class specification (“`class`”), and a connections dictionary (“`connections`”) specifying the chain (“`chain`”), terminus direction (“`direction`”), and residue positions that were available for fusion (“`residues`”) were included in the json file. Geometric class specification for oligomeric scaffolds required an N or C either following the oligomeric architecture, which defined the terminus available for subsequent fusions. A connections  dictionary specifying each chain, terminus direction, and residue positions allowed for fusion was required for each chain and terminus direction. The oligomeric scaffold asymmetric unit only required one connections dictionary, specifying fusion to only one terminus and one chain. As monomers, helical repeat proteins require a list of two connections dictionaries as they allow fusions to either N or C terminus. An example json file example is shown below where the connections dictionary specifies allowable fusions on the C terminus of the pH trimer from residue 50 and over, and anywhere along the helical repeat proteins:

```
{
{"file": "/path/to/ph_trimer.pdb",
  "class": ["C3_C"],
```

```
  "connections":[
    {"chain": 1, "direction": "C", "residues":["50:"]}
 ]
},
{"file": "/path/to/helicalrepeatprotein1.pdb",
  "class": ["Monomer"],
  "connections": [
   {"chain": 1, "direction": "N", "residues":[":210"]},
   {"chain": 1, "direction": "C", "residues":["-210:"]}
  ]
},
{"file": "/path/to/helicalrepeatprotein2.pdb",
  "class": ["Monomer"],
  "connections": [
   {"chain": 1, "direction": "N", "residues":[":160"]},
   {"chain": 1, "direction": "C", "residues":["-160:"]}
  ]
}
}
```

To prepare the WORMS software for fusions, we also included a map for backbone connections (`--bbconn`), which specifies the order for which scaffolds of each class could be fused. We used the following backbone connection map, which specifies that the N terminus of the fusion of two helical repeat proteins in the monomer class be fused to the C terminus of an oligomer with C3 symmetry:

```
--bbconn C3 C3_C    \
         NC Monomer \
         N_ Monomer \
```

Finally, we specified a `--geometry NullCriteria()` option, which allowed generating arbitrary fusions of varying radius and length to the C terminus of the pH trimer.

#### Computational docking of plugged antibody nanoparticles

The generated fusions served as docking inputs into the antibody nanoparticle through the (`--inputs1` and `--inputs2` options). To prepare the antibody nanoparticle for axle docking along its C3 axis, we generated a context structure pdb file by isolating the chains forming the C3 symmetric aperture. We fed this context structure and each trimer fusion separately into the RPXDock application [3] using the axle docking protocol (`--architecture AXLE_C3`). To enable trimming, we set a maximum number of trimmed residues (`--max_trim`) and specified the chains representing the fusion to be trimmed (`--trimmable_components`). Finally, to enable greater sampling, we allowed the RPXDock application to sample the entire search space (`--docking_method`) at a cartesian and angular resolution of 0.25 angstroms and degrees, respectively (`--grid_resolution_cart_angstroms` and `--grid_resolution_ori_degrees`). An example axle docking executable is shown below.

```
PYTHONPATH=/path/to/rpxdock/site-packages /path/to/python/environment
/path/to/rpxdock/dock.py \
                --architecture AXLE_C3 \
                --inputs1 /path/to/plugfusion.pdb \
                --inputs2 /path/to/contextstructure.pdb \
                --max_trim 400 \
                --cart_bounds -300 300 \
                --docking_method grid \
```

```
--grid_resolution_cart_angstroms 0.25 \
--grid_resolution_ori_degrees 0.25 \
--hscore_files ailv_h \
--hscore_data_dir /path/to/rpxdock/hscore \
--trimmable_components “A” \
--function stnd \
```

We evaluated the docked interfaces between each fusion and antibody nanoparticle using the RPX score and NContact metrics, which are predictive of geometric complementarity and designability. Empirically, we deemed docks with a RPX score greater than 50 and greater than one residue pair contact to be suitable for sequence optimization.

#### Computational design of plugged antibody nanoparticles

We used the Rosetta Macromolecular Modeling suite to optimize the sequence residues contributing to either the fusion domains or interface between plug and nanoparticle. With residue selectors, we identified clashing residue positions as a result of the WORMS fusion and residues contributing to the interface between the plug and antibody nanoparticle. The sequence of the selected residues were optimized based on the local positions of each residue defined by secondary structure, solvent accessible surface area (SASA) of the backbone and of the Cβ atom, and number of amino acid side chains in a cone extending along the CA-CB vector side chain neighbors. An example of the fusion, docking, and design of O432-17 is included at [https://github.com/erincyang/plug_design.git](https://github.com/erincyang/plug_design.git).

#### Small scale bicistronic bacterial protein expression for trimeric plug and tetrameric designs for sfGFP-Fc lysate assembly

Synthetic genes bicistronically encoding designed trimeric plug and tetramer sequences were purchased from Genscript in pET29b+ vectors and connected by an intergenic region (‘TAAAGAAGGAGATATCATATG') and a C-terminal 6xHistidine tag on the tetramer. Expression plasmids were transformed into BL21(DE3) *e. Coli* cells and grown in LB medium supplemented with 50 mg/L Kanamycin at 37°C overnight. The overnight culture was diluted into autoinduction media and incubated 37°C overnight. Cells were lysed chemically in BugBuster supplemented with 1mM PMSF and 20 mM imidazole, and cleared by centrifugation. Clarified lysates were incubated with 3uM final concentration of sfGFP-Fc and purified by immobilized metal affinity chromatography (IMAC) with Ni-NTA magnetic beads. The soluble fractions were washed with 25 mM Tris pH 8.0, 300 mM NaCl, 60 mM imidazole before eluting with 25 mM Tris pH 8.0, 300 mM NaCl, 300 mM Imidazole. Elution fractions from bicistronic expression plasmids were subsequently subjected to Native PAGE and SDS-PAGE to identify slow migrating species that coeluted three proteins of different molecular weights indicating assembly of three-component species.

#### Production of Fc and Fc-fusions

Fc, sfGFP-Fc, and Cetuximab IgG were cloned into CMVR and transfected into in ExpiHEK293F cells and purified by IMAC with 50 mM Tris pH8.0, 300 mM NaCl, 500 mM imidazole elution buffer and purified further via size exclusion chromatography over a Superdex 200 10/300 GL FPLC column (Cytiva) into 50 mM Tris, pH 8.0, 300 mM NaCl, 0.75% CHAPS (Fc) or 50 mM Tris, pH 8.0, 300 mM NaCl, 0.05% glycerol (Cetuximab IgG, sfGFP-Fc). Stocks were frozen at -80 for subsequent analyses.

#### Large scale expression and purification of O432-17 components

Designs appearing to co-purify and yielding slowly migrating species by native PAGE were subsequently subcloned into pet29b+ vectors each encoding either a trimeric plug or a tetramer variant–both with C terminal hexahistidine tags–and expressed at larger scale (1 to 12 liters of culture). Cells were lysed by microfluidization in 25 mM Tris pH 8.0, 300 mM NaCl, 1 mM DTT, 1mM PMSF, 0.1mg/ml DNAse and cleared by centrifugation. Clarified lysates were filtered through 0.7um filters and purified by IMAC via gravity columns with nickel-NTA resin or HisTrap HP columns (Cytiva) using 25 mM Tris pH 8.0, 300 mM NaCl, 60 mM Imidazole wash buffer and 25 mM Tris pH 8.0, 300 mM NaCl, 300 mM Imidazole elution buffer. Elution fractions containing pure proteins were concentrated using centrifugal filter devices (Millipore) and further purified on a Superdex 200 10/300 GL (for large scale purification) or Superose 6 10/300 GL (for comparison to assembly) gel filtration column (Cytiva) using 25 mM Tris pH 8.0, 150 mM NaCl, 0.75% CHAPS. Gel filtration fractions containing pure protein in the desired oligomeric state were pooled, concentrated and frozen in aliquots at -80°C for subsequent analyses.

#### In vitro assembly of O432-17 nanoparticles

Purified tetramer and Fc or IgG from gel filtration fractions were assembled in a 1:1 molar ratio and with 1.1x excess trimeric plug. Molar ratios were calculated based on the monomeric extinction coefficient. Assemblies were assembled between 100-500 uL between 5-50 uM, dialyzed overnight at 25°C into 25 mM Tris pH 8.0, 150 mM NaCl, and purified on a Superose 6 10/300 GL gel filtration column (Cytiva) using 25 mM Tris pH 8.0, 150 mM NaCl as the running buffer.

In vitro assembly of O432-17 nanoparticles with molecular cargoes were assembled in the same molar ratio of purified tetramer, Fc or Cetuximab IgG, and trimeric plug from gel filtration fractions with 1x purified pos36GFP per monomer of component or 3x pegRNA per nanoparticle to conserve RNA material. Nanoparticles were screened for in vitro packaging by SEC on a Superose 6 10/300 GL column (Cytiva) in either low or high salt Tris buffer (25 mM Tris, 200 mM NaCl pH 8.0, 25 mM Tris, 1 M NaCl, pH 8.0).

#### Dynamic light scattering

DLS measurements were performed using the default Sizing and Polydispersity method on the UNcle (Unchained Labs). O432-17 variants (8.8 µl) were pipetted into the provided glass cuvettes. DLS measurements were run in triplicate at 25°C with an incubation time of 1 s; results were averaged across 10 runs and plotted using Python3.7.

#### Negative Stain Electron Microscopy preparation and data collection of O432-17 nanoparticles and variants

Pre- or post-SEC O432-17 assemblies between 0.1 and 0.2 mg/ml in 25 mM Tris pH 8.0, 150 mM NaCl were applied onto 400- or 200-mesh carbon-coated copper grids and glow discharged for 20 s, followed by 3x application of 3.04 µl 2% nano-W or Uranyless stain.
Micrographs were recorded using EPU software (Thermo Fisher) on a 120 kV Talos L120C transmission electron microscopy (Thermo Fisher) at 45,000 nominal magnification (pixel size: 3.156 Å per pixel) at a defocus range of 1.0 to 2.5 µm.

#### Negative Stain Electron Microscopy data analysis of O432-17 nanoparticles and variants

Negative Stain Electron Microscopy datasets were processed by Relion3.0 software [4]. Micrographs were imported into the Relion3.0 software and a contrast transfer function was estimated using GCTF [5]. Around 500 particles were manually picked, 3D classified, and selected classes were used as templates for particle picking in all images. Approximately 100k picked particles were 2D classified for 25 iterations into 50 classes.

#### Cryo-Electron Microscopy preparation and data collection of O432-17 nanoparticles

2 ul of pre-SEC purified O432-17-Fc sample at 0.5 mg/ml in 25 mM Tris pH 8.0, 150 mM NaCl was applied onto C-flat 1.2/1.3 holey carbon grids. Grids were then plunge-frozen into liquid ethane and cooled with liquid nitrogen using a ThermoFisher Vitrobot Mk IV with 6.5-s blotting time and 0 blot force. The blotting process took place inside the vitrobot chamber at 22°C and 100% humidity. Data acquisition was performed with Leginon on a Titan Krios electron microscope operating at 300 kV using a K3 summit direct electron detector equipped with an energy filter and operating in super-resolution mode. The nominal magnification for data collection was 105000X with a calculated pixel size of 0.42 Å/pixel, with a final dose of 63.775 $e^{-}/A^{2}$ for 2223 movies.

#### Cryo-Electron Microscopy data analysis of O432-17 nanoparticles

All data processing was carried out in CryoSPARC v3.0.0 [6]. Alignment of movie frames was performed using Patch Motion with an estimated B-factor of 500 $Å^2$, with a maximum alignment resolution set to 5. During alignment, all movies were fourier cropped by ½. Defocus and astigmatism values were estimated using Patch CTF with default parameters. 1,391 nanoparticle particles were initially manually picked and extracted with a box size of 576 pixels. This was followed by a round of 2D classification and subsequent template-picking using the best 2D class averages low-pass filtered to 20 Å. Particles were next picked with Template Picker and were manually inspected before extracting with a box size of 660 pixels, and further fourier cropped to a final box size of 330 pixels, for a total of 66,904 particles. A round of reference-free 2D classification was next performed in CryoSPARC with a maximum alignment resolution of 6 Å. The best classes which revealed visibly assembled nanoparticles were used for 3D ab initio determination using the C1 symmetry operator. This was followed by a round of 3D heterogeneous refinement using C1 symmetry and sorting into 4 distinct classes, all of which revealed complete plugging of the octahedral 3-component nanoparticle. Thus, all 50,017 of the best particles selected from 2D classification were subjected to non-uniform 3D refinement with octahedral symmetry applied, yielding a final map with an estimated global resolution of 7.06 Å, following per-particle defocus refinement. The final maps were deposited in the EMDB under accession number EMD-29602. Relaxed models were generated by rigid-body docking followed by relaxing the design model into the final cryo-EM density map in Rosetta [7].

#### Computational design of O432-17 electrostatically charged variants

A consensus design approach was used to first identify interior surface positions predicted to be the most robust to surface mutations. These positions were divided into three tiers based on the predicted enhancement to stability and / solubility. Using the Rosetta modeling suite, the trimeric plug design model was redesigned by allowing optimization of the identities of interior surface residues that did not contribute to the interface between plug monomers or between the trimeric plug and antibody nanoparticle. We deployed three tiers of sequence optimization strategies per charged variant. The first strategy enabled optimization of residue positions toward only charged amino acid identities (arginine or lysine for the positively charged variants, glutamate and aspartate for the negatively charged variants). The second strategy included the charged amino acid identities but also polar, non-charged amino acid identities such as asparagine, glutamine, and alanine to maintain helical propensity. The third strategy included the residue identities in the first and second strategy, but also enabled reversion back to the original identity in the trimeric plug design model. Mutations that resulted in losses of significant atomic packing interactions or side chain-side chain or side chain-backbone hydrogen bonds were discarded.

The best scoring design for each design strategy and surface position tier were selected for inclusion as variant proteins.

#### Expression and purification of electrostatically charged trimeric plug variants

Plasmids encoding electrostatically charged trimeric plug variants and containing a C terminal hexahistidine tag were cloned into pet29b+ vectors and expressed overnight at 37°C in autoinduction media [8]. Cells were lysed by microfluidization or sonication in 25 mM Tris pH 8.0, 500 mM NaCl, 1 mM DTT, 1mM PMSF, 0.1mg/ml DNAse and cleared by centrifugation. Clarified lysates were filtered through 0.7um filters and purified by IMAC via gravity columns with nickel-NTA resin or HisTrap HP columns (Cytiva) using 25 mM Tris pH 8.0, 500 mM NaCl, 60 mM Imidazole wash buffer and 25 mM Tris pH 8.0, 500 mM NaCl, 300 mM Imidazole elution buffer. Elution fractions containing pure proteins were concentrated using centrifugal filter devices (Millipore) and further purified on a Superdex 200 10/300 GL (for large scale purification) or Superose 6 10/300 GL (for comparison to assembly) gel filtration column (Cytiva) using 25 mM Tris pH 8.0, 500 mM NaCl, 0.75% CHAPS. Gel filtration fractions containing pure protein in the desired oligomeric state were pooled, concentrated and frozen in aliquots at -80°C for subsequent analyses.

#### Gel electrophoresis

Native agarose gels were prepared using 0.8% ultrapure agarose (Invitrogen) in TAE buffer (Thermo Fisher) containing SYBR Gold (Invitrogen). 9 µL of O432-17 nucleocapsids were treated with 1 µL of Benzonase (Invitrogen) at 25 °C for 30 min, followed by mixing with 2uL 6x loading dye (NEB, no SDS), and electrophoresed at 120 V for 30 min. Gels were imaged for RNA and subsequently stained by Gelcode Blue for protein (ThermoFisher).

Protein SDS-PAGE gels were performed using anyKD polyacrylamide gels (Bio-Rad) in Tris-glycine buffer.

#### Rational design of O432-17 pH-responsive variants

Bulky hydrophobic residues such as isoleucine and leucine amino acid positions within the pH-responsive trimeric interface were selected for optimization to either alanine or valine using the Rosetta Software Suite. The point mutation with the best scoring interface energy (ddG) at each position was selected as a trimeric plug variant. Combinatorial pairs of each point mutation variant were also included as trimeric plug variants.

#### AF647 conjugation to O432-17 trimeric plug variants

Trimeric plug variants containing a T359C mutation were generated by site directed mutagenesis PCR. Alexa Fluor™ 647 C2 Maleimide (Thermo Fisher Scientific) dissolved in DMSO and trimeric plug variants containing 10x TCEP were incubated in a 5:1 molar ratio with respect to the trimeric plug monomer for 16 hours (overnight) at 4°C in PBS + 0.75% CHAPS titrated to pH 7.2. The maleimide reaction was quenched with 1mM DTT and buffer exchanged into 25 mM Tris pH 8.0, 150 mM NaCl using PD-10 desalting columns with Sephadex-25 resin (Cytiva) according to manufacturer’s protocol and dialyzed for 3-4 days in 25 mM Tris pH 8.0, 150 mM NaCl, 0.75% CHAPS at 4°C. Positively charged trimeric plug variants were buffer exchanged and dialyzed into 25 mM Tris pH 8.0, 500 mM NaCl, 0.75% CHAPS.

#### Flow-cytometry based pH-titration

Linear myc peptide with an N terminal lysine side chain and 3x glycine linker (KGGGEQKLISEEDL) was produced via solid-phase peptide synthesis and biotinylated via amide formation. The resulting biotinylated myc peptide was purified by RP-HPLC and quality checked for the proper molecular weight via LC-MS, lyophilized, and dissolved in 100% DMSO for long term storage. 3.0-3.4 μm streptavidin coated polystyrene particles (Spherotech) were incubated with biotinylated myc peptide diluted in PBS to 5% DMSO for 20 minutes at 25°C. Following two washes in PBS + 3% BSA by pelletting at 3000G for 5 minutes, coated polystyrene particles were incubated with α-myc-O432-17 nanoparticle variants for one hour at 25°C. The coated particles were washed in PBS + 3% BSA twice and split equally into pH-titrated citrate-phosphate buffers for 30 minutes at 25°C. Coated particles were washed twice with PBS + 3% BSA and resuspended for flow-cytometry.

All flow-cytometry experiments were performed on a LSR II Flow Cytometer (BD Biosciences). Lasers were calibrated with coated particles stained with either FITC Anti-Myc tag antibody (Abcam 9E10), APC Anti-Myc tag antibody (Abcam 9E10), or no antibody. 10,000 events were collected per sample on three biological replicates. All flow-cytometry results were analyzed using the FlowJo, LLC software. Normalization to the minimum and maximum fluorescence of each channel with each titration sample was performed in Python3.7, and the apparent pKa of AF647 and sfGFP fluorescence was estimated with a four parameter nonlinear logistic regression fit in GraphPad Prism version 9.3.1 for Windows, GraphPad Software, San Diego, California USA, www.graphpad.com.

#### Cells

WT HeLa (ATCC CCL-2), EGFR KO HeLa (Abcam ab255385), and A431 cells (ATCC CRL-1555) were cultured at 37 °C with 5% $CO_2$ in flasks with Dulbecco's modified Eagle medium (DMEM) (Gibco) supplemented with 1 mM L-glutamine (Gibco), 4.5 g/liter D-glucose (Gibco), 10% fetal bovine serum (FBS) (Hyclone) and 1% penicillin-streptomycin (PenStrep) (Gibco). WT HeLa and EGFR KO HeLa also cultured with 1× nonessential amino acids (Gibco) supplemented into the media. Cells were passaged twice per week. To passage, cells were dissociated using 0.05% trypsin EDTA (Gibco) and split 1:5 or 1:10 into a new tissue culture (TC)–treated T75 flask (Thermo Scientific ref 156499).

#### Immunostaining

35 mm glass bottom dishes seeded at a density of 20k cells / dish. A final monomeric concentration of 10 nM of O432-17-CTX or O432-17-Fc nanoparticles were incubated with cultured cells in serum-free DMEM. Cells were fixed 4% paraformaldehyde, permeabilized with 100% methanol, and blocked with PBS + 1% BSA. Cells were immunostained with Anti-LAMP2A antibody (Abcam ab18528) followed by goat anti-rabbit- IgG Alexa Fluor™ 488 secondary antibody (Thermo Fisher A-11034) and 4′,6-diamidino-2-phenylindole (DAPI) (Thermo Fisher D1306) and stored in the dark at 4°C until imaging.

#### Nanoparticle uptake confocal microscopy (A431 and HeLa EGFR KO Cells)

Cells were washed twice with FluoroBrite DMEM imaging media and subsequently imaged in the same media in the dark at room temperature. Epifluorescence imaging was performed on a Yokogawa CSU-X1 spinning dish confocal microscope with either a Lumencor Celesta light engine with 7 laser lines (408, 445, 473, 518, 545, 635, 750 nm) or a Nikon LUN-F XL laser launch with 4 solid state lasers (405, 488, 561, 640 nm), 40x/0.95 NA objective and a Hamamatsu ORCA-Fusion scientific CMOS camera, both controlled by NIS Elements software (Nikon). The following excitation/emission filter combinations (center/bandwidth in nm) were used: BFP: EX408, EM443/38, GFP: EX473 EM525/36, RFP: EX545 EM605/52, Far Red: EX635 EM705/72. Exposure times were 100 ms for the acceptor direct channel and 500ms for all other channels, with no EM gain set and no ND filter added. All epifluorescence experiments were subsequently analyzed using Image J.

#### Nanoparticle uptake image acquisition (HeLa WT Cells)

4-color, 3D images were acquired with a commercial OMX-SR system (GE Healthcare). Toptica diode lasers with excitation at 405nm, 488nm, and 640nm were used. Emission was collected on three separate PCO.edge sCMOS cameras using an Olympus 60× 1.42NA PlanApochromat oil immersion lens. 1024×1024 images (pixel size 6.5 μm) were captured with no binning. Acquisition was controlled with AcquireSR Acquisition control software. Z-stacks were collected with a step size of 125 nm. Images were deconvolved in SoftWoRx 7.0.0 (GE Healthcare) using the enhanced ratio method and 200 nm noise filtering. Images from different color channels were registered in SoftWoRx using parameters generated from a gold grid registration slide (GE Healthcare).

### Supplementary Figures

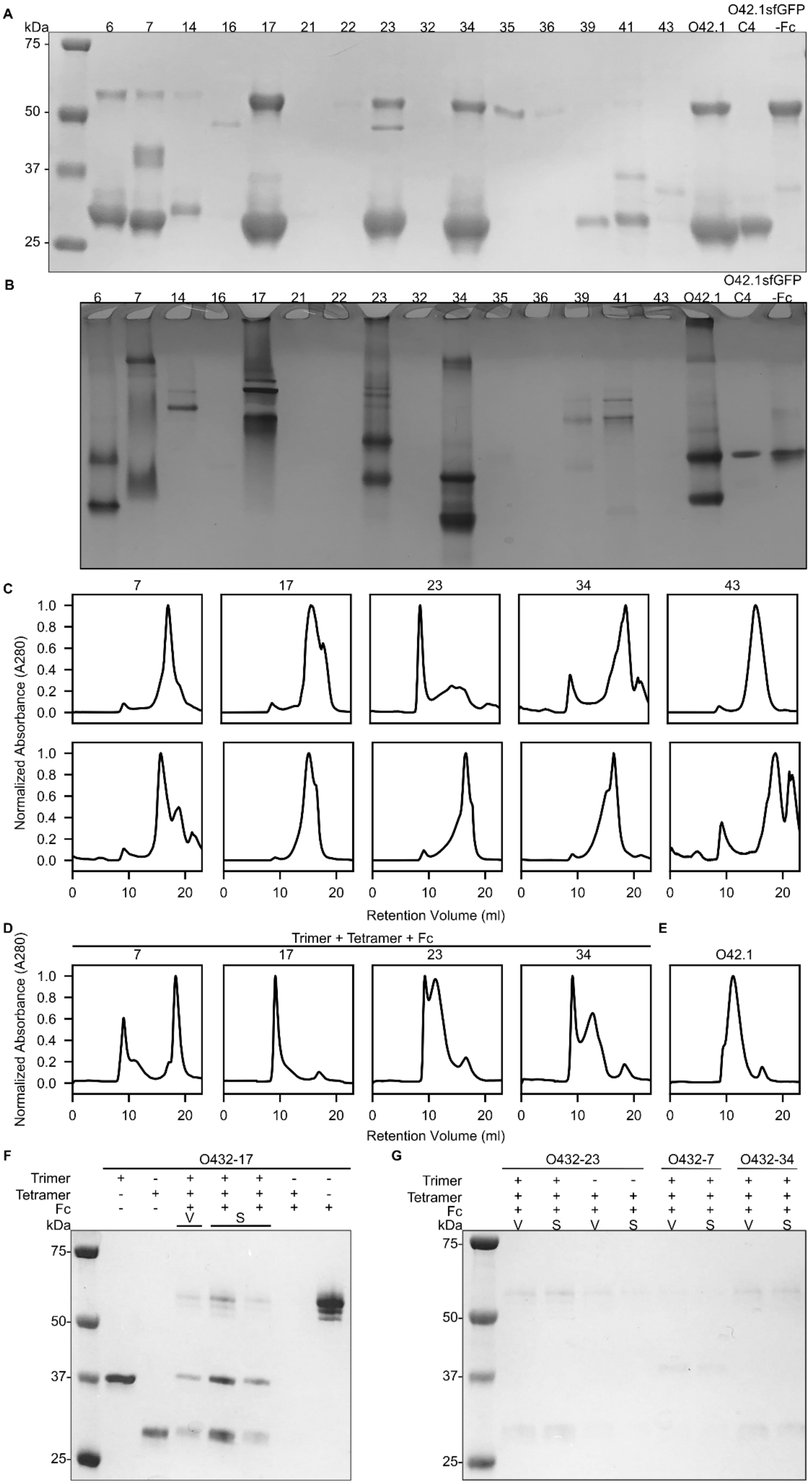
A
kDa
6 7 14 16 17 21 22 23 32 34 35 36 39 41 43 O42.1 C4 O42.1sfGFP -Fc
75
50
37
25
B
6 7 14 16 17 21 22 23 32 34 35 36 39 41 43 O42.1 C4 O42.1sfGFP -Fc
C
7 17 23 34 43
Trimer
Tetramer
Normalized Absorbance (A280)
Retention Volume (ml)
D
Trimer + Tetramer + Fc
7 17 23 34
E
O42.1
Normalized Absorbance (A280)
Retention Volume (ml)
F
O432-17
Trimer
Tetramer
Fc
kDa
V
S
75
50
37
25
G
O432-23
O432-7
O432-34
Trimer
Tetramer
Fc
kDa
75
50
37
25

**Figure S1: Example experimental screen of O432 nanoparticles. A.** Clarified lysates of 16 designs where the designed trimer co-eluted with the tetramer were supplemented with a purified sfGFP-Fc fusion protein, purified by IMAC, and subject to reducing SDS-PAGE, **B.** followed by non-denaturing native PAGE gel electrophoresis. All designs were compared against the previously designed, two-component antibody nanoparticle (O42.1) [9]**,** O42.1 tetrameric component and sfGFP-Fc. **C.** Subcloned components from putative three-component assemblies were purified separately, **D.** and, material permitting, after stoichiometric *in vitro* assembly by SEC. **E.** Non-reducing SDS-PAGE of the void (V) and shoulder (S) peaks from SEC of O432-17 with purified O432-17 tetramer + Fc assembly and trimer, tetramer, and Fc components as controls, and **F.** of the void (V) and shoulder (S) peaks from SEC purification of O432 assemblies.

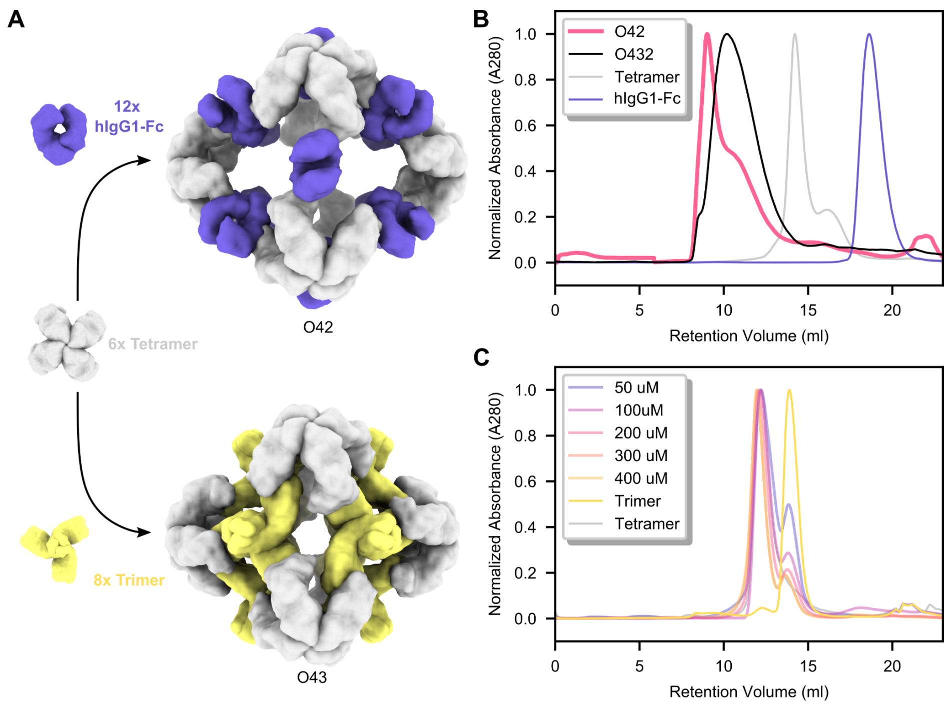


**Figure S2: Mixing two of the three individual components does not result in nanoparticle assembly. A.** 6× designed tetramers form protein-protein interfaces with both 12× Fc and 8× designed trimers. A schematic depicts a hypothetical nanoparticle assembly from the designed tetramer with only the Fc (top) or only the trimer (bottom). **B.** Representative SEC traces on the Superose 6 10/300 GL of assembly reactions containing only the designed tetramer and Fc (pink), compared to the full three-component assembly (black) and the individual components (gray and purple). **C.** Representative SEC traces on the Superdex 200 10/300 GL of assembly reactions containing only the designed tetramer and trimer at concentrations of 50 μM, 100 μM, 200 μM, 300 μM, and 400 μM, compared to the individual components (gray and yellow).

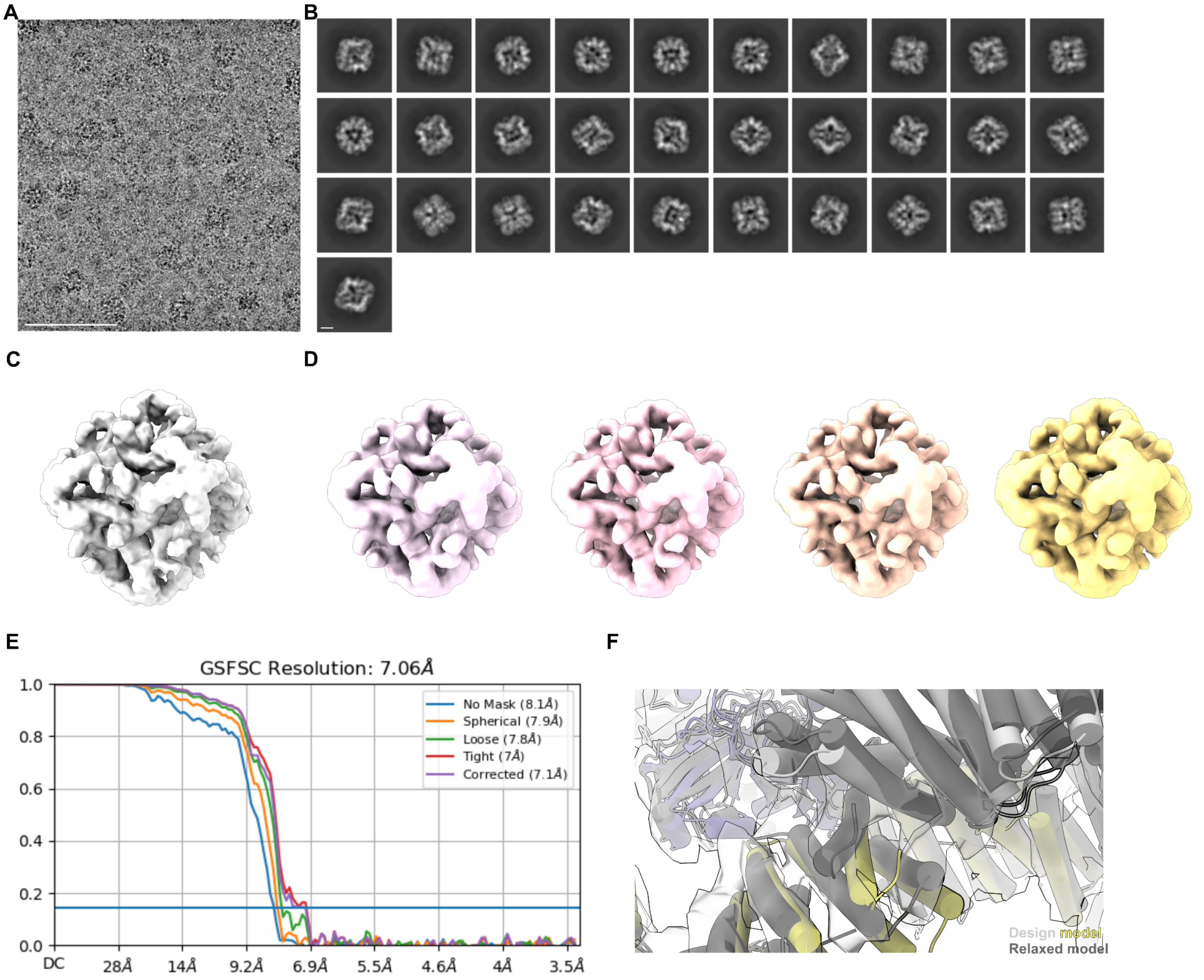


**Figure S3: Cryo-EM processing shows newly designed trimeric plug occupying all 3-fold symmetry axes of the nanoparticle octahedral architecture. A.** Representative micrograph of cryo-EM sample. Scale bar, 100 nm. **B.** Reference-free two-dimensional class averages. Scale bar, 10 nm. **C.** *Ab initio* three-dimensional reconstruction without applied octahedral symmetry. **D.** 3D reconstructions generated following a heterogeneous refinement in the absence of applied octahedral symmetry. All four classes show trimeric plugs occupying all facets of the designed nanoparticle. **E.** Gold-standard Fourier shell correlation curves for the O432-Fc EM density map with octahedral symmetry applied. **F.** Close up of plug interface between design model (light gray and yellow) and model built from the 3D reconstruction (dark gray).

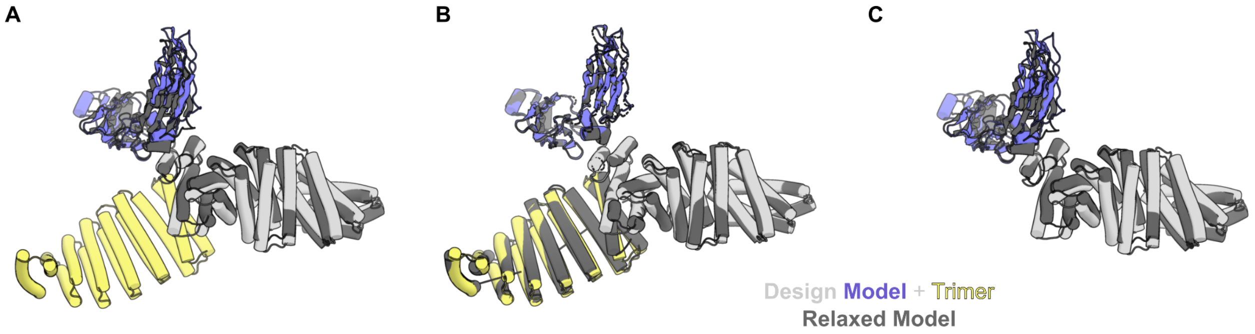


**Figure S4: Comparison of design models of O42.1 and O432-17 with models relaxed into Cryo-EM density. A.** The three-component O432-17 design model (light gray, purple, and yellow) overlaid on the two-component O42.1 relaxed model (dark gray) shows an RMSD of 1.9 Å. **B.** The O432-17 design model (light gray, purple, and yellow) deviates from its relaxed model (dark gray) by 1.6 Å. **C.** The O42.1 design model deviates from its relaxed model (dark gray) by 4.2 Å.

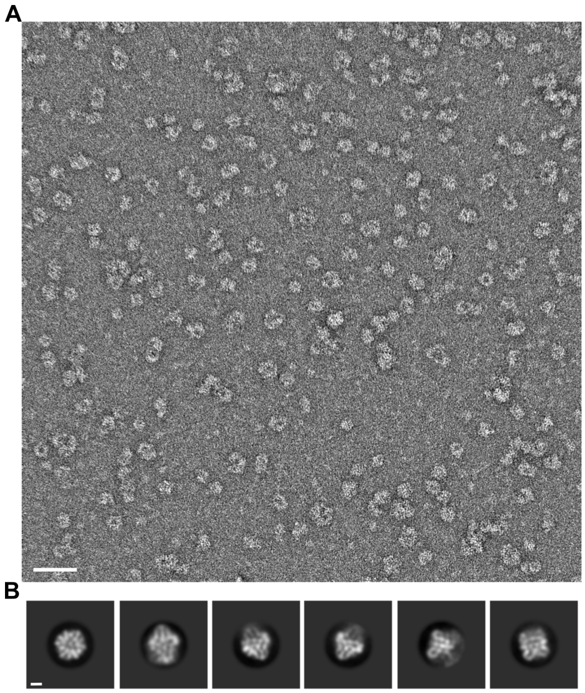

**Figure S5: Negative stain electron microscopy of O432-17 nanoparticle assembly in the presence of RNA. A.** Representative negatively stained electron micrograph of O432-17(+) in the presence of RNA. Scale bar, 100 nm. **B.** Reference-free two-dimensional class averages showing multiple views of O432-17 nanoparticles. Scale bar, 10 nm.

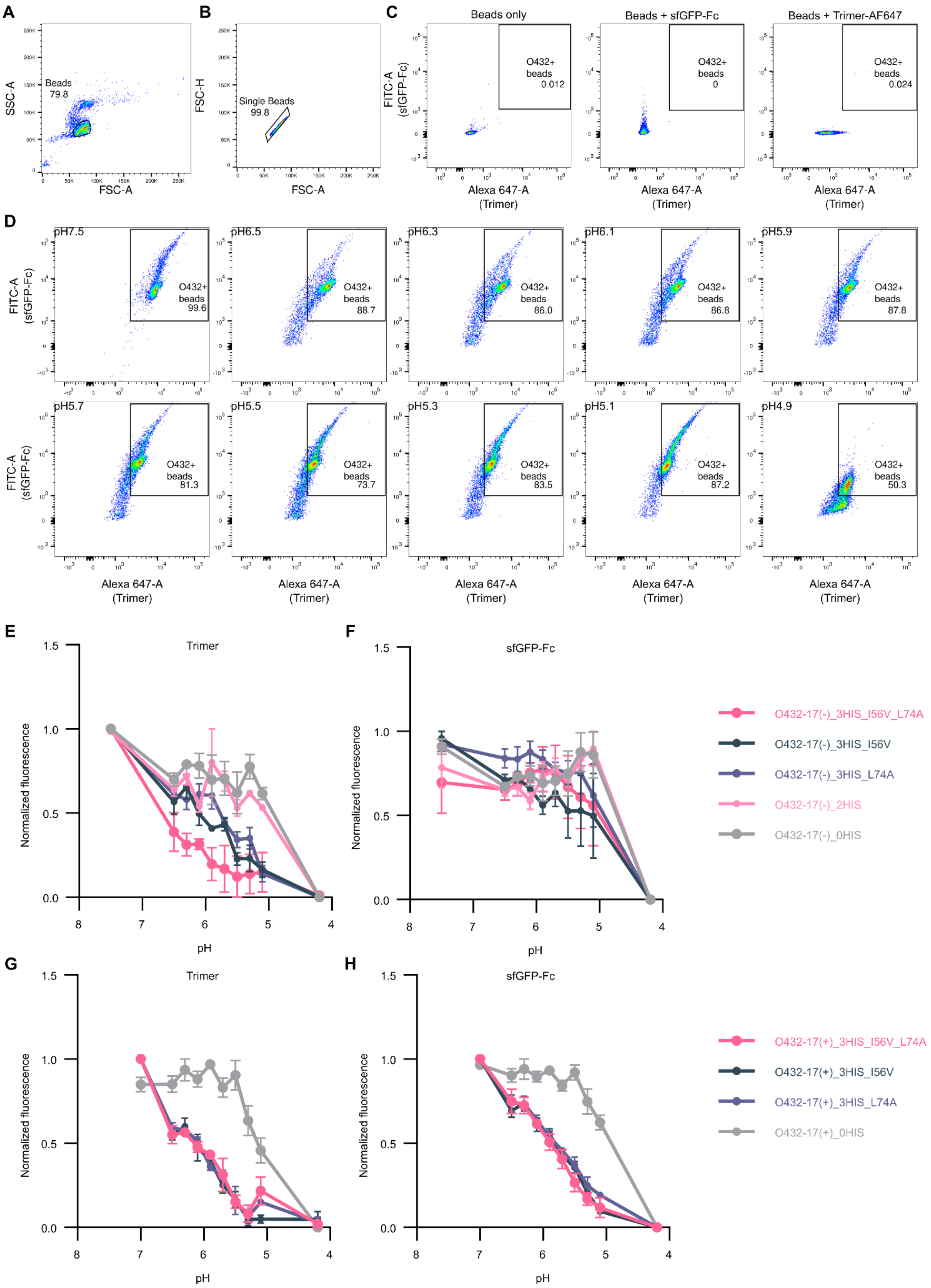
A
SSC-A
FSC-A
Beads
79.8
B
FSC-H
FSC-A
Single Beads
99.8
C
Beads only
Beads + sfGFP-Fc
Beads + Trimer-AF647
FITC-A (sfGFP-Fc)
O432+ beads 0.012
O432+ beads 0
O432+ beads 0.024
Alexa 647-A (Trimer)
D
pH7.5
pH6.5
pH6.3
pH6.1
pH5.9
O432+ beads 99.6
O432+ beads 88.7
O432+ beads 86.0
O432+ beads 86.8
O432+ beads 87.8
pH5.7
pH5.5
pH5.3
pH5.1
pH4.9
O432+ beads 81.3
O432+ beads 73.7
O432+ beads 83.5
O432+ beads 87.2
O432+ beads 50.3
FITC-A (sfGFP-Fc)
Alexa 647-A (Trimer)
E
Trimer
F
sfGFP-Fc
Normalized fluorescence
pH
O432-17(-)_3HIS_I56V_L74A
O432-17(-)_3HIS_I56V
O432-17(-)_3HIS_L74A
O432-17(-)_2HIS
O432-17(-)_0HIS
G
Trimer
H
sfGFP-Fc
Normalized fluorescence
pH
O432-17(+)_3HIS_I56V_L74A
O432-17(+)_3HIS_I56V
O432-17(+)_3HIS_L74A
O432-17(+)_0HIS

**Figure S6: Flow cytometry analysis of O432-17(-) and O432-17(+) nanoparticle variants. A.** Forward and side scattering intensity isolated all beads. **B.** Isolated beads were selected for singlet scattering. **C.** Singlet beads were gated for both AF647 and sfGFP-Fc signal with reference to negative controls: beads only (left), beads incubated with sfGFP-Fc (middle), and beads incubated with AF647-labeled trimeric plug (right). **D.** The mean fluorescence intensity of beads positive for trimer and sfGFP-Fc signal was taken at each pH. This gating strategy was applied across all trimeric plug variants: 0HIS, 2HIS, 3HIS_I56V, 3HIS_L74A, 3HIS_I56V_L74A. Mean fluorescence intensity of O432-17(-) nanoparticles was measured as a function of pH for the **E.** trimeric plug variants and **F.** sfGFP-Fc. Mean fluorescence intensity of O432-17(+) nanoparticles was measured as a function of pH for the **G.** trimeric plug variants and **H.** sfGFP-Fc.

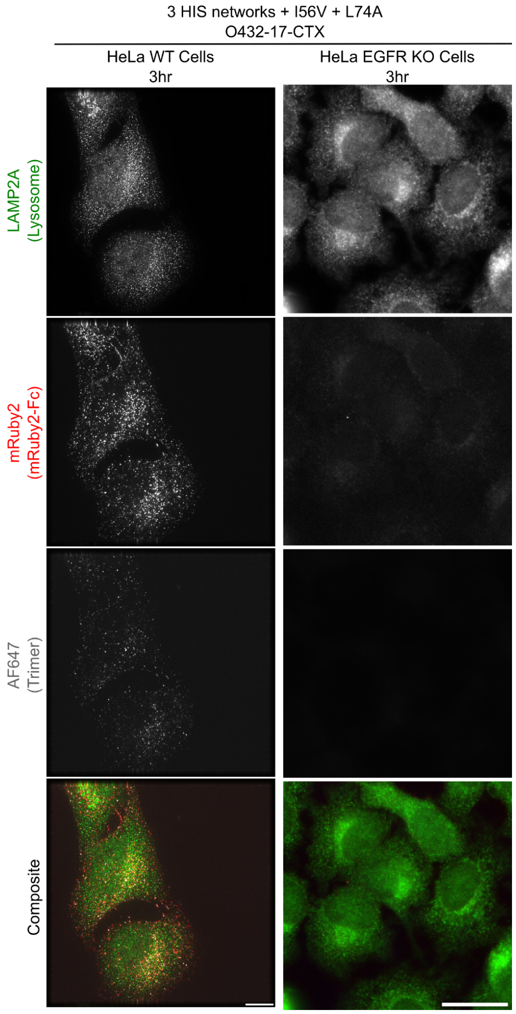

**Figure S7: Targeted receptor-mediated uptake of O432-17-CTX nanoparticles in WT and EGFR KO HeLa cells after 3 hours.** Scale bar, 10 μm.

### Supplementary Tables

##### Table 1: Cryo-EM data collection and refinement statistics

| | O432-17 |
|---|---|
| Microscope | Krios |
| Voltage (kV) | 300 |
| Detector | Gatan K3 Summit |
| Recording mode | Super-resolution |
| Magnification | 105,000 X |
| Movie micrograph pixel size (Å) | 0.42 |
| Dose rate ($e^{-}$/Å$^2$/s) | 20 |
| No. of frames per movie micrograph | 75 |
| Frame exposure time (ms) | 40 |
| Movie micrograph exposure time (s) | 3 |
| Total dose ($e^{-}$/Å$^2$) | 63.775 |
| Under focus range (μm) | 0.8 - 1.7 |
| Number of movie micrographs | 2,223 |
| Total number of picked particles | 88,357 |
| Particles in the final reconstruction | 50,017 |
| Map symmetry | O |
| Map resolution (GS-FSC) | 7.05 Å |
| B-factor | -230.0 |
| EMDB ID | EMD-29602 |

##### Table 2: Amino acid sequences of assembling O432-17 designs

| **Design name** | **Sequence** |
|---|---|
| O432-17-C3 | MSEEKIEKLLEELTASTAELKRATASLRAITEELKKNPSEDALVEHNRAIVEHNAIIVENNRIIATVLLAIVAAIATNEATLAADKAKEAGASEVAKLAKKVLEEAEELAKENDSEEALKVVKAIADAAKAAAEAAREGKTEVAKLALKVLEEAIELAKENRSEEALKVVREIARAALAAAQAAEEGKTEVAKLALKVLEEAIELAKENRSEEALKVVREIARAAL |

| | |
|---|---|
| | AAAQAAEEEGKTEVAKLALEVLEQAIEAAKLQRSERALEMVREIARAALEAARN<br>AEGGRSDRARAILASLKVSIIVVKLKSSGTSEEEILRIVLKIIKELRKTAKESGQSAS<br>YIATMEAEIVKAIDYALDLSGTSGSWSGLEHHHHHH |
| O432-17-C4 | MFNKDQQSAFYEILNMPNLNEALRNGFIQLLKDDPSKSEVILTAALIAAKLSEDIR<br>TLKESGSSYEEIAERVARAVALLVALLKTNGVSEDEIALAVALIISAVIQTLKESGSS<br>YEVIAEIVARIVAEIVEALKRSGTSEDEIAEIVARVISEVIRTLKESGSSYEVIAEIVAR<br>IVAEIVEALKRSGTSEDEIAKIVARVIAEVLRTLKESGSSEEVIKEIVARIITEIKEALK<br>RSGTSEDEIELITLMIEAALEIAKLKSSGSEYEEIAEDVARRIAELVEKLKRDGTSA<br>VEIAKIVAAIISAVIAMLKASGSSYEVIAEIVARIVAEIVEALKRSGTSAAIIALIVALVIS<br>EVIRTLKESGSSFEVILEIVIRIVLEIIEALKRSGTSEQDVMLIVMAVLLVVLATLQLS<br>GSGSWSGLEHHHHHH |
| Negatively charged variants | |
| O432-17(-)_2HIS-C3 | MSEEKIEKLLEELTASTAELKRATASLRAITEELKKNPSEDALVEHNRAIVEHNAIIV<br>ENNRIIATVLLAIVAAIATNEATLAADKAKEAGASEVAELAKEVLEEAEELAKENDS<br>EEALKVVKAIADAAKAAAEAAREGKTEVAELALKVLEEAIELAKENRSEEALKVV<br>REIARAALAAAQAAEEGKTEVAELALEVLEEAIELAKENRSEEALKVVREIARAAL<br>AAAQAAEEGKTEVAELALEVLEQAIEAAKLQRSERALEMVREIARAALEAARNA<br>EGGRSDRARAILASLKVSIIVVKLKSSGTSEEEILRIVLKIIKELRKEAKEEGQSAS<br>YIATMEAEIVKAIDYALDLSGTSGSWSGLEHHHHHH* |
| O432-17(-)_3HIS_I56V_L74A-C3 | MMSEEKIEKLLEELTAATAELKRATASLRAITEELKKNPSEDALVEHNRAIVEHNAI<br>VVEHNRIIATVLLAIVAAAATNEATLAADKAKEAGASEVAELAKEVLEEAEELAKE<br>NDSEEALKVVKAIADAAKAAAEAAREGKTEVAELALKVLEEAIELAKENRSEEAL<br>KVVREIARAALAAAQAAEEGKTEVAELALEVLEEAIELAKENRSEEALKVVREIAR<br>AALAAAQAAEEGKTEVAELALEVLEQAIEAAKLQRSERALEMVREIARAALEAAR<br>NAEGGRSDRARAILASLKVSIIVVKLKSSGTSEEEILRIVLKIIKELRKEAKEEGQS<br>ASYIATMEAEIVKAIDYALDLSGCSGSWSGLEHHHHHH |
| O432-17(-)_0HIS-C3 | MMSEEKIEKLLEELTASTAELKRSTASLRASTEELKKNPSEDALVENNRLIVENNA<br>IIVENNRIIATVLLAIVAAIATNEATLAADKAKEAGASEVAELAKEVLEEAEELAKEN<br>DSEEALKVVKAIADAAKAAAEAAREGKTEVAELALKVLEEAIELAKENRSEEALK<br>VVREIARAALAAAQAAEEGKTEVAELALEVLEEAIELAKENRSEEALKVVREIARA<br>ALAAAQAAEEGKTEVAELALEVLEQAIEAAKLQRSERALEMVREIARAALEAARN<br>AEGGRSDRARAILASLKVSIIVVKLKSSGTSEEEILRIVLKIIKELRKEAKEEGQSA<br>SYIATMEAEIVKAIDYALDLSGCSGSWSGLEHHHHHH |
| O432-17(-)_3HIS_I56V-C3 | MSEEKIEKLLEELTAATAELKRATASLRAITEELKKNPSEDALVEHNRAIVEHNAIV<br>VEHNRIIATVLLAIVAAIATNEATLAADKAKEAGASEVAELAKEVLEEAEELAKEND<br>SEEALKVVKAIADAAKAAAEAAREGKTEVAELALKVLEEAIELAKENRSEEALKV<br>VREIARAALAAAQAAEEGKTEVAELALEVLEEAIELAKENRSEEALKVVREIARAA<br>LAAAQAAEEGKTEVAELALEVLEQAIEAAKLQRSERALEMVREIARAALEAARNA<br>EGGRSDRARAILASLKVSIIVVKLKSSGTSEEEILRIVLKIIKELRKEAKEEGQSAS<br>YIATMEAEIVKAIDYALDLSGCSGSWSGLEHHHHHH |
| O432-17(-)_3HIS_L74A-C3 | MSEEKIEKLLEELTAATAELKRATASLRAITEELKKNPSEDALVEHNRAIVEHNAIIV<br>EHNRIIATVLLAIVAAAATNEATLAADKAKEAGASEVAELAKEVLEEAEELAKEND<br>SEEALKVVKAIADAAKAAAEAAREGKTEVAELALKVLEEAIELAKENRSEEALKV<br>VREIARAALAAAQAAEEGKTEVAELALEVLEEAIELAKENRSEEALKVVREIARAA<br>LAAAQAAEEGKTEVAELALEVLEQAIEAAKLQRSERALEMVREIARAALEAARNA |

| | |
|---|---|
| | EGGRSDRARAILASLKVSIIVVKLKSSGTSEEEILRIVLKIIKELRKEAKEEGQSAS YIATMEAEIVKAIDYALDLSGCSGSWSGLEHHHHHH* |
| Positively charged variants | |
| O432-17(+)_2HIS-C3 | MSEEKIEKLLEELTASTAELKRATASLRAITEELKKNPSEDALVEHNRAIVEHNAIIV ENNRIIATVLLAIVAAIATNEATLAADKAKEAGASEVAKLAKKVLKQAEQLAKENDS EEALKVVKAIADAAKAAAEAAREGKTEVAKLALKVLANAIKLAKENRSEEALKVV REIARAALAAAQAAEEGKTEVARLALKVLQNAIQLAKENRSEEALKVVREIARAA LAAAQAAEEGKTEVAKRALKVLQQAIQAAKLQRSERALEMVREIARAALEAARN AEGGRSDRARAILASLQVSIIVVKLKSSGTSEEEILRKVLKIIKELRKKAKEQGQS ASYIATMEAEIVKAIDYALDLSGTSGSWSGLEHHHHHH* |
| O432-17(+)_3HIS_I56V-C3 | MSEEKIEKLLEELTAATAELKRATASLRAITEELKKNPSEDALVEHNRAIVEHNAIV VEHNRIIATVLLAIVAAIATNEATLAADKAKEAGASEVAKLAKKVLKQAEQLAKEND SEEALKVVKAIADAAKAAAEAAREGKTEVAKLALKVLANAIKLAKENRSEEALKV VREIARAALAAAQAAEEGKTEVARLALKVLQNAIQLAKENRSEEALKVVREIARA ALAAAQAAEEGKTEVAKRALKVLQQAIQAAKLQRSERALEMVREIARAALEAAR NAEGGRSDRARAILASLQVSIIVVKLKSSGTSEEEILRKVLKIIKELRKKAKEQGQ SASYIATMEAEIVKAIDYALDLSGCSGSWSGLEHHHHHH |
| O432-17(+)_3HIS_L74A-C3 | MSEEKIEKLLEELTAATAELKRATASLRAITEELKKNPSEDALVEHNRAIVEHNAIIV EHNRIIATVLLAIVAAAATNEATLAADKAKEAGASEVAKLAKKVLKQAEQLAKEND SEEALKVVKAIADAAKAAAEAAREGKTEVAKLALKVLANAIKLAKENRSEEALKV VREIARAALAAAQAAEEGKTEVARLALKVLQNAIQLAKENRSEEALKVVREIARA ALAAAQAAEEGKTEVAKRALKVLQQAIQAAKLQRSERALEMVREIARAALEAAR NAEGGRSDRARAILASLQVSIIVVKLKSSGTSEEEILRKVLKIIKELRKKAKEQGQ SASYIATMEAEIVKAIDYALDLSGCSGSWSGLEHHHHHH |
| O432-17(+)_0HIS-C3 | MSEEKIEKLLEELTASTAELKRSTASLRASTEELKKNPSEDALVENNRLIVENNAII VENNRIIATVLLAIVAAIATNEATLAADKAKEAGASEVAKLAKKVLKQAEQLAKEND SEEALKVVKAIADAAKAAAEAAREGKTEVAKLALKVLANAIKLAKENRSEEALKV VREIARAALAAAQAAEEGKTEVARLALKVLQNAIQLAKENRSEEALKVVREIARA ALAAAQAAEEGKTEVAKRALKVLQQAIQAAKLQRSERALEMVREIARAALEAAR NAEGGRSDRARAILASLQVSIIVVKLKSSGTSEEEILRKVLKIIKELRKKAKEQGQ SASYIATMEAEIVKAIDYALDLSGCSGSWSGLEHHHHHH |

**Table 3: Amino acid sequences of hIgG1-Fc, Fc-fusions, and IgGs produced and used to assemble O432-17 designs**

| Design name | Sequence |
|---|---|
| hIgG1-Fc | EPKSSDKTHTCPPCPAPELLGGPSVFLFPPKPKDTLMISRTPEVTCVVVDVSHE DPEVKFNWYVDGVEVHNAKTKPREEQYNSTYRVVSVLTVLHQDWLNGKEYKC KVSNKALPAPIEKTISKAKGQPREPQVYTLPPSRDELTKNQVSLTCLVKGFYPSDI AVEWESNGQPENNYKTTPPVLDSDGSFFLYSKLTVDKSRWQQGNVFSCSVMH EALHNHYTQKSLSLSPGK |

| | |
|---|---|
| sfGFP-Fc | SRATMETDTLLLWVLLLWVPGSTGHHHHHHGGSENLYFQGGSSKGEELFTGVVPILVELDGDVNGHKFSVRGEGEGDATNGKLTLKFICTTGKLPVPWPTLVTTLTYGVQCFSRYPDHMKRHDFFKSAMPEGYVQERTISFKDDGTYKTRAEVKFEGDTLVNRIELKGIDFKEDGNILGHKLEYNFNSHNVYITADKQKNGIKANFKIRHNVEDGSVQLADHYQQNTPIGDGPVLLPDNHYLSTQSVLSKDPNEKRDHMVLLEFVTAAGITHGMDELYKGGSGSEPKSSDKTHTCPPCPAPELLGGPSVFLFPPKPKDTLMISRTPEVTCVVVDVSHEDPEVKFNWYVDGVEVHNAKTKPREEQYNSTYRVVSVLTVLHQDWLNGKEYKCKVSNKALPAPIEKTISKAKGQPREPQVYTLPPSRDELTKNQVSLTCLVKGFYPSDIAVEWESNGQPENNYKTTPPVLDSDGSFFLYSKLTVDKSRWQQGNVFSCSVMHEALHNHYTQKSLSLSPGK |
| mRuby2-Fc | SRATMETDTLLLWVLLLWVPGSTGHHHHHHGGSENLYFQGGSVSKGEELIKENMRMKVVMEGSVNGHQFKCTGEGEGNPYMGTQTMRIKVIEGGPLPFAFDILATSFMYGSRTFIKYPKGIPDFFKQSFPEGFTWERVTRYEDGGVVTVMQDTSLEDGCLVYHVQVRGVNFPSNGPVMQKKTKGWEPNTEMMYPADGGLRGYTHMALKVDGGGHLSCSFVTTYRSKKTVGNIKMPGIHAVDHRLERLEESDNEMFVVQREHAVAKFAGLGGGMDELYKGGSGSEPKSSDKTHTCPPCPAPELLGGPSVFLFPPKPKDTLMISRTPEVTCVVVDVSHEDPEVKFNWYVDGVEVHNAKTKPREEQYNSTYRVVSVLTVLHQDWLNGKEYKCKVSNKALPAPIEKTISKAKGQPREPQVYTLPPSRDELTKNQVSLTCLVKGFYPSDIAVEWESNGQPENNYKTTPPVLDSDGSFFLYSKLTVDKSRWQQGNVFSCSVMHEALHNHYTQKSLSLSPGK |
| Cetuximab light chain | MELGLSWIFLLAILKGVQCDILLTQSPVILSVSPGERVSFSCRASQSIGTNIHWYQQRTNGSPRLLIKYASESISGIPSRFSGSGSGTDFTLSINSVESEDIADYYCQQNNNWPTTFGAGTKLELKRTVAAPSVFIFPPSDEQLKSGTASVVCLLNNFYPREAKVQWKVDNALQSGNSQESVTEQDSKDSTYSLSSTLTLSKADYEKHKVYACEVTHQGLSSPVTKSFNRGEC |
| Cetuximab heavy chain | MELGLSWIFLLAILKGVQCQVQLKQSGPGLVQPSQSLSITCTVSGFSLTNYGVHWVRQSPGKGLEWLGVIWSGGNTDYNTPFTSRLSINKDNSKSQVFFKMNSLQSNDTAIYYCARALTYYDYEFAYWGQGTLVTVSAASTKGPSVFPLAPSSKSTSGGTAALGCLVKDYFPEPVTVSWNSGALTSGVHTFPAVLQSSGLYSLSSVVTVPSSSLGTQTYICNVNHKPSNTKVDKRVEPKSCDKTHTCPPCPAPELLGGPSVFLFPPKPKDTLMISRTPEVTCVVVDVSHEDPEVKFNWYVDGVEVHNAKTKPREEQYNSTYRVVSVLTVLHQDWLNGKEYKCKVSNKALPAPIEKTISKAKGQPREPQVYTLPPSREEMTKNQVSLTCLVKGFYPSDIAVEWESNGQPENNYKTTPPVLDSDGSFFLYSKLTVDKSRWQQGNVFSCSVMHEALHNHYTQKSLSLSPGKGSGHHHHHH |

**Table 4: Amino acid sequences of molecular cargoes used by O432-17 designs**

| **Design name** | **Sequence** |
|---|---|
| pos36-GFP | MGHHHHHHGGASKGERLFRGKVPILVELKGDVNGHKFSVRGKGKGDATRGKLTLKFICTTGKLPVPWPTLVTTLTYGVQCFSRYPKHMKRHDFFKSAMPKGYVQERTISFKKDGKYKTRAEVKFEGRTLVNRIKLKGRDFKEKGNILGHKLRYNFNSHKVYITADKRKNGIKAKFKIRHNVKDGSVQLADHYQQNTPIGRGPVLLPRNHYLSTRSKLSKDPKEKRDHMVLLEFVTAAGIKHGRDERYK |

| | |
|---|---|
| pegRNA | mC*mC*mA*rGrGrCrUrUrCrCrGrGrGrUrCrArUrCrCrCrGrUrUrUrArGrArGrCrUrArGrArArArUrArGrCrArArGrUrUrArArArArUrArArGrGrCrUrArGrUrCrCrGrUrUrArUrCrArArCrUrUrGrArArArArArGrUrGrGrCrArCrCrGrArGrUrCrGrGrUrGrCrGrCrArCrCrUrGrGrUrGrUrArUrGrArCrCrCrGrGrArCrGrCrGrGrUrUrCrUrArUrCrUrArGrUrUrArCrGrCrGrUrUrArArArCrCrArArCrUrA*mG*mA*mA |

**Table 5: Details on EM data acquisition on different O432-17 samples**

| Sample name | Stain | Magnification | Pixel size (Å/pixel) | # Micrographs |
|---|---|---|---|---|
| O432-17 Fc | Uranyless | 45,000 | 3.156 | 289 |
| O432-17 CTX | Uranyless | 45,000 | 3.156 | 249 |
| O432-17(+) RNA CTX | Uranyless | 45,000 | 3.156 | 169 |

**Table 6: Details on EM data processing on different O432-17 samples**

| Sample name | Particle picking | CTF estimation | 2D class averages | # particles in final selected 2D classes/total picked particles |
|---|---|---|---|---|
| O432-17 Fc | CisTEM | CTFFIND4 within Relion | Relion | 84107/112334 |
| O432-17 CTX | Relion template picking | CTFFIND4 within Relion | Relion | 23676/84107 |
| O432-17(+) RNA CTX | CryoSPARC template picking | CTFFIND4 within CryoSPARC | CryoSPARC | 18204/79682 |

**Table 7: Statistical information for pH titration experiments.** All analyses were performed using Graphpad Prism version 9.3.1 Software.

| Experiment (Fig.) | Dunnett's multiple comparisons test | Mean Diff. | 95.00% CI of diff. | Below threshold ? | Summary | Adjusted P Value |
|---|---|---|---|---|---|---|
| Fig 4G/S6D | O432-17(-)_0HIS vs. O432-17(-)_2HIS | 0.06277 | -0.2448 to 0.3703 | No | ns | 0.9598 |
| | O432-17(-)_0HIS vs. O432-17(-)_3HIS_I56V | 0.25 | -0.05761 to 0.5575 | No | ns | 0.1407 |

| | | | | | | |
|---|---|---|---|---|---|---|
| | O432-17(-)_0HIS vs. O432-17(-)_3HIS_I56V_L74A | 0.388 | 0.08047 to 0.6956 | Yes | ** | 0.0092 |
| | O432-17(-)_0HIS vs. O432-17(-)_3HIS_74A | 0.1996 | -0.1080 to 0.5071 | No | ns | 0.3019 |
| Fig 4H/S6E | O432-17(-)_0HIS vs. O432-17(-)_2HIS | 0.02994 | -0.2523 to 0.3122 | No | ns | 0.9964 |
| | O432-17(-)_0HIS vs. O432-17(-)_3HIS_I56V | 0.1138 | -0.1684 to 0.3960 | No | ns | 0.6984 |
| | O432-17(-)_0HIS vs. O432-17(-)_3HIS_I56V_L74A | 0.07698 | -0.2052 to 0.3592 | No | ns | 0.8962 |
| | O432-17(-)_0HIS vs. O432-17(-)_3HIS_L74A | -0.02741 | -0.3096 to 0.2548 | No | ns | 0.9974 |
| Fig S6F | O432-17(+)_0HIS vs. O432-17(+)_3HIS_I56V | 0.3731 | 0.04387 to 0.7024 | Yes | * | 0.0231 |
| | O432-17(+)_0HIS vs. O432-17(+)_3HIS_I56V_L74A | 0.3502 | 0.02095 to 0.6794 | Yes | * | 0.0349 |
| | O432-17(+)_0HIS vs. O432-17(+)_3HIS_L74A | 0.3564 | 0.02720 to 0.6857 | Yes | * | 0.0312 |
| Fig S6G | O432-17(+)_0HIS vs. O432-17(+)_3HIS_I56V | 0.3094 | -0.02647 to 0.6454 | No | ns | 0.0766 |
| | O432-17(+)_0HIS vs. O432-17(+)_3HIS_I56V_L74A | 0.3221 | -0.01386 to 0.6580 | No | ns | 0.0627 |
| | O432-17(+)_0HIS vs. O432-17(+)_3HIS_L74A | 0.2852 | -0.05070 to 0.6211 | No | ns | 0.1109 |
